# Type III Druantia two-component antiphage defense depends on the DruH-DruE interaction for halting phage DNA cyclization and replication

**DOI:** 10.64898/2026.05.17.725784

**Authors:** Yakun Li, Zheng-Guo He

## Abstract

Bacteria have evolved multiple immune systems to resist phage invasion, however, only a small part of the defensive mechanisms have been clearly uncovered. In this study, we report a type III Druantia two-component defense system, DruH-DruE, identified from *Mycobacterium smegmatis*. The DruH-DruE prevents phage DNA cyclization and replication.DruE can be replaced from the defense system by either homolog in *M. tuberculosis* or *M. smegmatis*. The physical interaction between this two components is essential for fighting against phage infection. Mutations in the interaction sites led to the loss of phage-defending function of the system. The broad-spectrum antiphage ability of the defense system could be activated by the small tail protein Gp25 of phage A10ZJ24. This study fills a major gap in current knowledge of antiphage mechanism of type III Druantia defense system, expanding our understanding of the immune mechanisms in prokaryotic cells.

## Introduction

In the arms race between bacteriophages or phages and hosts, bacteria have evolved multiple immune systems to resist phage invasion, and these systems are usually clustered as the defense islands in the genome (1, 2). So far, only a small part of the defense mechanisms of these innate immune systems have been uncovered. For example, the CBASS system (Cyclic oligonucleotide based anti-phage signaling systems) is able to sense viral RNA to produce circular dinucleotides and activate downstream effector proteins to trigger abortion infection (Abi) (3, 4). The nuclease helicase immunity (Nhi) can block phage DNA accumulation (5). The Hachiman defense system is activated to degrade all DNA in the cell causing abortive infection (6). In addition, some other prokaryotic antiviral mechanisms have been gradually elucidated, including BREX (Bacteriophage Exclusion) (7), Sir2-HerA (8–10), pAgos (prokaryotic Argonautes) (11–12), DISARM (Defence Island System Associated with Restriction-Modification) (13), RADAR (Restriction by an Adenosine Deaminase acting on RNA) (14–15), Gabija (16, 17) and Shedu (18, 19). These studies have greatly expanded our understanding of the immune mechanisms in prokaryotic cells.

Helicase is a common component of multiple eukaryotic immune system (20). Interestingly, helicases in prokaryotes are also involved in many antiviral processes (7, 8, 13, 17, 21). For example, the helicase proteins of the BREX (BrxHI) system, Hachiman (HamB) system and hna system contain a DExD/H box helicase domain, which is crucial for combating phage infection (6, 21). There are also some DExD/H box helicase proteins reported in mycobacteria (22–24), but it is currently unclear whether these helicases function during phage infection.

The Druantia system is a potential antiviral defense system characterized by the Sorek team, which can be divided into three types. Type I consists of five genes called DruABCDE. Type II is composed of four genes called DruMFGE, while type III is composed of DruH and DruE genes (1). These three Druantia systems all contain a highly conserved DruE protein, but there is no recognized domains in the remaining component genes. The DruE protein contains an unknown functional domain (DUF1998), a helicase feature, and a Walker A/B motif that suggests ATP utilization. The DUF1998 domain is one of the components of genes in multiple defense systems (1, 13, 25). Although the antiphage mechanisms of many prokaryotic defense systems have been elucidated in recent years, there have been few studies on the Druantia system, only demonstrating its antiphage activity or synergistic antiphage activity with other defense systems (1, 26, 27), but its specific mechanism is still unclear.

Mycobacteria belong to a class of important actinomycetes, including pathogenic *Mycobacterium tuberculosis* and *Mycobacterium abscesses* (28, 29), as well as non-pathogenic and fast-growing *Mycobacterium smegmatis* (30). Recently, some defense systems in *M. smegmatis* have also been found to play important roles in resisting phage invasion, such as the BREX-like system (31). However, it is believed that there are some unidentified defense systems in mycobacteria.

In this study, we identified a type III Druantia defense system in the genome of *M. smegmatis*, encoded by *MSMEG_1253-MSMEG_1254*. These two genes are highly conserved in *Actinobacteria*. We provide evidence to show that the type III Druantia is a two-component antiphage defense system. Both DruH and DruE are essential for combating phage infection, and the defense function of the Druantia system also depends on the interaction between this two-components. Furthermore, the system was found to be activated by the small tail protein Gp25 of phage A10ZJ24. Our study uncovered the antiphage mechanism of type III Druantia system, providing important insight into the prokaryotic antiviral defense.

## Results

### The type III Druantia system of *M. smegmatis* confers antiphage activity

According to the previous report (1), we noticed that the type III two-component Druantia defense system is widely present in *Actinobacteria* (Figure. 1A), containing two conserved ORFs, DruH and DruE. Sequence blast analysis of different DruH or DruE homologs showed that the type III Druantia defense system exhibits high conservation at the amino acid residue level (Figure S1 and 2). However, the function of DruH is not yet clear. The DruE protein is annotated as a DExD/H box helicase, which includes three domains: DExD/H box (pfam00271), helicase C-terminal (pfam00271), and DUF1998 (Figure S3A). To confirm the potential antiviral function of these conserved DruH-DruE systems in *Actinobacteria*, we first cloned the complete DNA sequence fragment encoding the predicted type III Druantia defense system in *M. smegmatis*, including the 500 bp upstream of the start codon of the *druH* gene until to the stop codon of the *druE* gene. The recombinant vector was then transformed into *M. smegmatis* for expressing the DruHE system (Figure 1B). As shown in the Figure 1C, when series of mycobacterial phages covering different groups were used to attack transformed strains, the recombinant DruH-DruE strains can provide protection to these widely diverse mycobacteriophages (Figure 1C and Figure S3B). Among them, the defense system exhibits strong resistance to A10ZJ24, A22GX2, A3GX4, and A9GX2, with phage efficiency of plating reduction of over 10^4^times (Figure 1C and Figure S3B).

**Figure 1.**
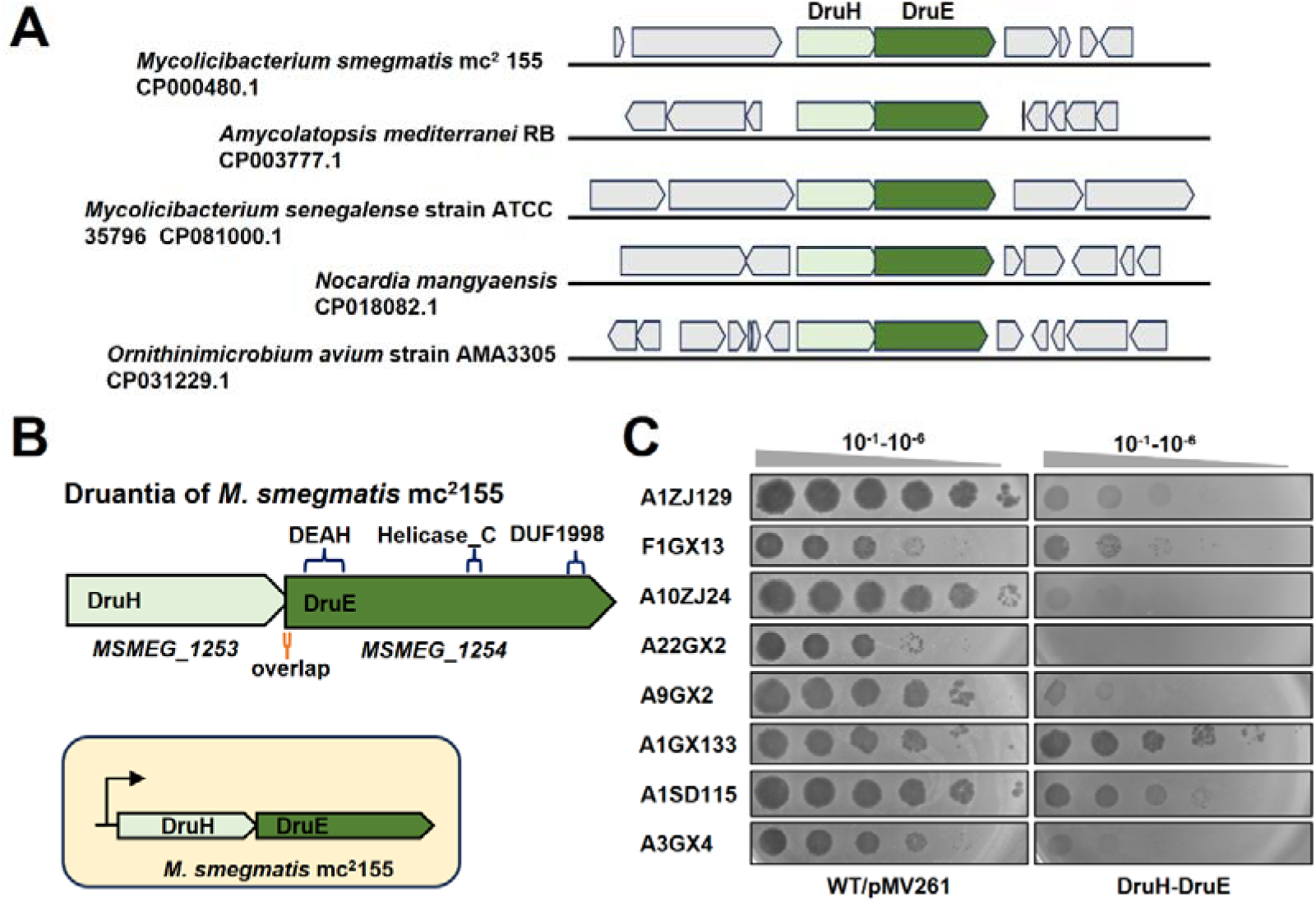
Type III Druantia system of *M. smegmatis* confers antiphage activity. (A) Conservation of the Type III Druantia Defense System in *Actinobacteria*. The type III Druantia defense system is widely conserved among *Actinobacteria*. In the schematic representation, the DruH gene is depicted in light green, while the DruE gene is shown in dark green. (B) Genetic Organization and Domain Architecture of the Type III Druantia System in *M. smegmatis* mc^2^155.The type III Druantia system in *M. smegmatis* mc^2^155 consists of two adjacent genes: *MSMEG_1253*, encoding the DruH protein (1,132 aa), and *MSMEG_1254*, encoding the DruE protein (1,667 aa), with a 17-bp overlap between the two genes. DruE harbors several conserved domains, including a DExD/H box domain (pfam00270), a helicase conserved C-terminal domain (pfam00271), and a DUF1998 domain (pfam09369). For functional analysis, the native Druantia system, including its promoter, was cloned and transformed into *M. smegmatis*. (C) Antiphage Activity of the Type III Druantia System. Plaque formation assays were performed to evaluate the antiphage activity of the type III Druantia system. Two *M. smegmatis* strains were tested: the wild-type strain (WT/pMV261) and the strain overexpressing the DruH-DruE. The phages used to assess the defense system’s activity are listed on the left. See also Figure S1, S2 and S3.

These results indicate that the conserved type III Druantia system encoded by *M. smegmatis* has a good antiphage function.

### Both DruH and DruE play important roles for antiphage defense

Next, we determined whether each component of the type III Druantia system is necessary for antiphage activity. We separately cloned the DruH and DruE genes together with their upstream 500 bp fragments, and then constructed their respective *M. smegmatis* expression strains. As shown in Figure S4A, expressing the DruE gene alone has no effect on the mycobacterial growth, but the expression of the DruH gene obviously inhibited the growth of *M. smegmatis*, even leading to bacterial death. Strikingly, the co-expression of DruH-DruE had no significant effect on the mycobacterial growth, thus the expression of DruE completely neutralized the lethal effect of DruH on the growth of mycobacterium. An ATc (Anhydrous Tetracycline) induction-dependent expression strain was then constructed for determining the potential effect of DruH on phage infection. As shown in Figure 4B, A10ZJ24 and A9GX2 phages can effectively infect *mycobacterium* strains, and normal plaque formation can be clearly observed with or without ATc induction for DruH gene expression. The resistant ability of the DruE-expressing strain to phage A10ZJ24 was reduced by approximately 100 times, but its resistance to phage A9GX2 was similar to that of the wild-type strain (Figure S4B, lower panel). Interestingly, compared with the effect of individual gene expression, DruH-DruE co-expression significantly enhanced the resistance of recombinant mycobacteria to phages, and its resistance to phage A10ZJ24 was reduced by at least 10^4^ times compared to wild type strains (Figure S4B, lower panel).

Next, we evaluated the importance of conserved amino acid residues for the antiviral function of DruH and DruE proteins. Based on the conservative analysis shown in Figure S1 and 2, we produced 7 point mutations on DruH including E1096/1099A, E842A, D240A, R536A, R864/865A, D1108A, and E998A. Six point mutations on DruE including K119A, D225A, A292D, A1016D, C1516A, and P800A were also performed. Then, these mutant genes were separately introduced into the gene cassette of DruH-DruE and their effects on phage defense function were examined. As shown in Figure 2, all the mutations mentioned above, except for DruH^E998A^-DruE and DruH-DruE^P800A^, greatly eliminated the antiviral function of the type III Druantia system, and the phage regained good ability to infect and lyse *mycobacterium* (Figure 2A, 2B and Figure S4C).

**Figure 2.**
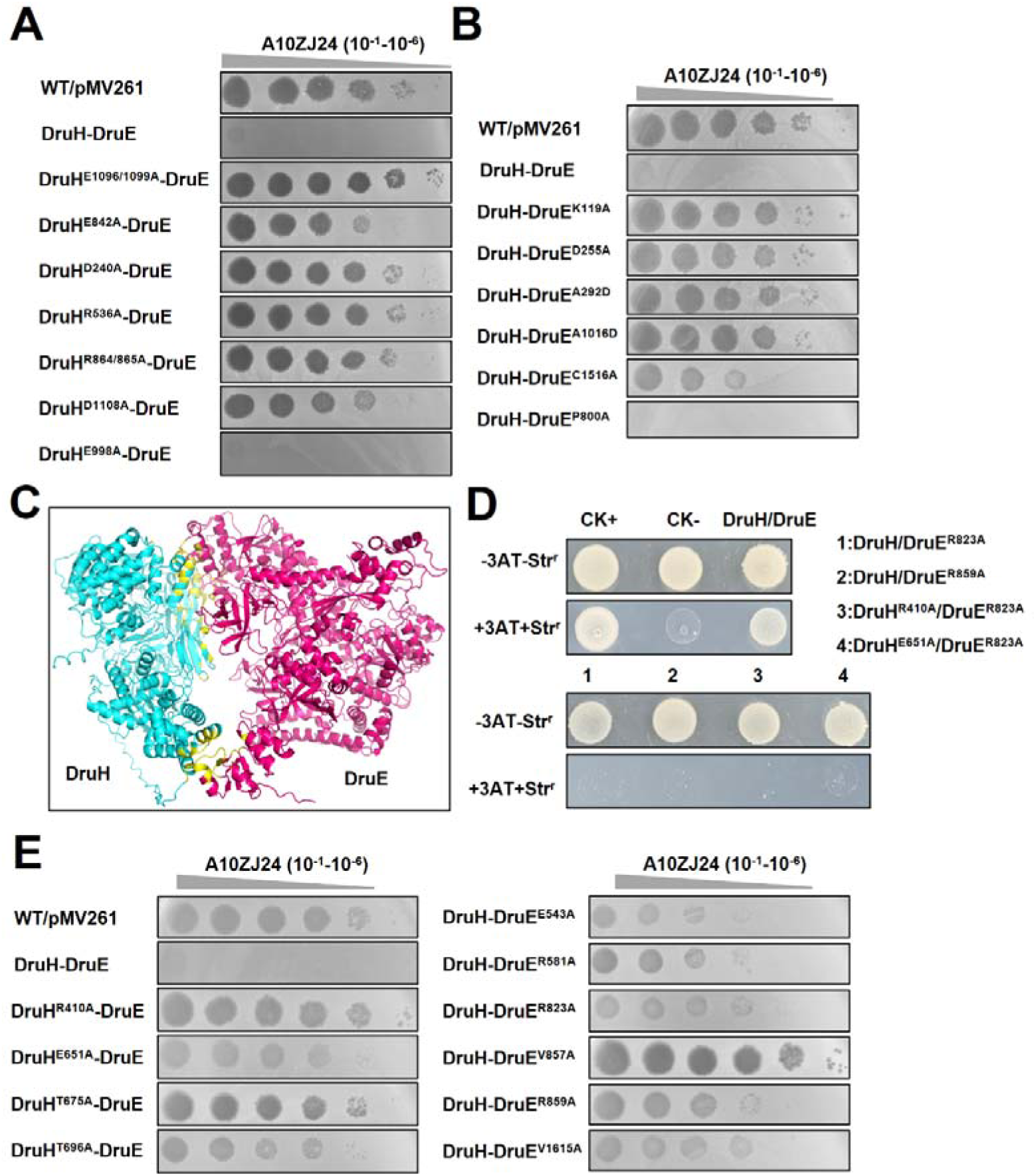
Both DruH and DruE are required for antiphage activity of the type III Druantia system. (A-B) Serial dilution assays for the plaque formation ability of phage A10ZJ24. Serial dilution assays were performed to assess the plaque-forming ability of phage A10ZJ24 on *M. smegmatis* strains. The wild-type strain (WT/pMV261) and the strain overexpressing the type III Druantia system (DruH-DruE) were used as controls.Additional strains included type III Druantia overexpression variants with point mutations at conserved sites in either DruH or DruE, enabling functional analysis of key residues. (C) Structural Prediction of the DruH-DruE Complex. The AlphaFold-predicted structure of the DruH-DruE complex reveals potential interaction regions. DruH is depicted in blue, DruE in red, and the putative interaction interface is highlighted in yellow. This structural model provides insights into the molecular basis of DruH-DruE binding and its role in antiphage activity. (D) Bacterial two-hybrid assays. Protein-protein interactions between DruH and DruE were validated using bacterial two-hybrid assays. *E. coli* reporter strains with various recombinant plasmids were spotted on the plate with or without streptomycin (str) and 3-amino-1, 2, 4-triazole (3-AT). Positive (CK^+^, pBT-*LGF2*/pTRG-*Gal11P*) and negative (CK^−^, pBT/pTRG) controls were included to confirm assay specificity. (E) Functional Impact of DruH-DruE Interface Mutations. The effect of point mutations at the DruH-DruE interaction interface on phage resistance was evaluated using plaque formation assays. The wild-type strain (WT/pMV261) and the strain overexpressing the type III Druantia system (DruH-DruE) were used as controls. Additional strains harboring mutations at the predicted DruH-DruE interface were tested to determine the functional importance of specific residues in the antiphage response. See also Figure S4.

These results indicate that both DruH and DruE components play important roles for the antiphage defense, and the conserved amino acid sites of DruH and DruE are crucial for the antiviral role of the defense system.

### DruH-DruE interaction is required for the antiphage defense function of type III Druantia system

To further determine the relationship between DruH and DruE, we used AlphaFold (32, 33) to predict the monomer structures of DruH and DruE (Figure S4D), and protein-protein interactions. The results showed that there may exist an interaction between DruH and DruE, and some key residues on the interaction interface of DruH-DruE, such as R410, R823, E651, R859, and V857 (Figure 2C and Figure S4C). Subsequently, we used bacterial two hybrid analysis to confirm the interaction between two proteins. As shown in Figure 2D, the negative control (CK^−^) strain did not grow on the screening medium supplemented with 3-AT. However, under the same experimental conditions, the DruH/DruE co-transformed strain containing wild type genes grew well on the screening medium and it was very similar to the positive interaction control strain (CK^+^), indicating a specific interaction between the two proteins encoded by DruH and DruE. Next, we performed site-directed mutagenesis on the residues of the DruH-DruE interface, and introduced these mutations into the DruH-DruE gene cassette. Thereafter, their interactions including DruH/DruE^R823A^, DruH/DruE^R859A^, DruH^R410A^/DruE^R823A^, and DruH^E651A^/DruE^R823A^, and their effect on phage defense function were determined. Significantly, these mutations resulted in a loss of growth ability of recombinant mycobacteria on the screening medium containing 3-AT (3-amino-1, 2, 4-triazole) (Figure 2D, bottom), indicating that there was no interaction between the proteins encoded by the mutant genes. Furthermore, we found that these predicted mutations at the DruH-DruE interface partially reduced or completely eliminated the antiviral function of the type III Druantia system (Figure 2C and Figure S4F).

Therefore, these results indicate that there exists an interaction between the proteins encoded by the *druH* and *druE* genes, and the interaction is required for the antiphage defense function of the III Druantia system.

### DruE homologs in different mycobacteria have similar defense function

Due to the difficulty in expressing and purifying DruE protein, we attempted to search for potential homologous genes in mycobacterial genomes. Interestingly, the proteins encoded by the *M. smegmatis* gene MSMEG_6160 and the *M. tuberculosis* gene MRA_3684 were found to have good conservation with the DruE protein. They both contain DExD/H box domains (pfam00271), helicase C-terminal domains (pfam00271), and DUF1998 domains, and the key amino acids are conserved (Figure 3A, top left and Figure S5A-S5C). Next, we examined whether MSMEG_6160 or MRA_3684 could replace the DruE protein in type III Druantia to exhibit the same activity. To this end, we introduced MSMEG_6160 or MRA_3684 into the DruH-DruE gene cassette to replace the DruE gene, and transformed it into a new recombinant strain of *M. smegmatis* (Figure 3A, upper right panel). Interestingly, consistent with the phenotype of DruH-DruE strains (Figure S4A), DruH-MRA_3684 and DruH-MSMEG_6160 strains effectively neutralized the toxic effects of DruH alone on mycobacterial cells (Figure S5D), and the recombinant strains showed significant resistance to phage A10ZJ24 and A9GX22 (Figure 3A, lower left panel and Figure S5E, lower left panel). Especially, the resistance to phage A10ZJ24 is strong, with efficiency of plating decrease of more than 10^4^ times, which is comparable to that of the DruH-DruE strain (Figure 1C). The resistance of DruH-MRA_3684 strain to phage A9GX2 is one order of magnitude stronger than that of DruH-MSMEGA-6160 strain (Figure S5E, lower left panel). Similar to DruE, the MRA_3684 gene alone or the MSMEG_6160 gene alone showed no defensive activity for the infection of either A10ZJ24 or A9GX2 phage (Figure 3A, lower right panel and Figure S5E, lower right panel).

**Figure 3.**
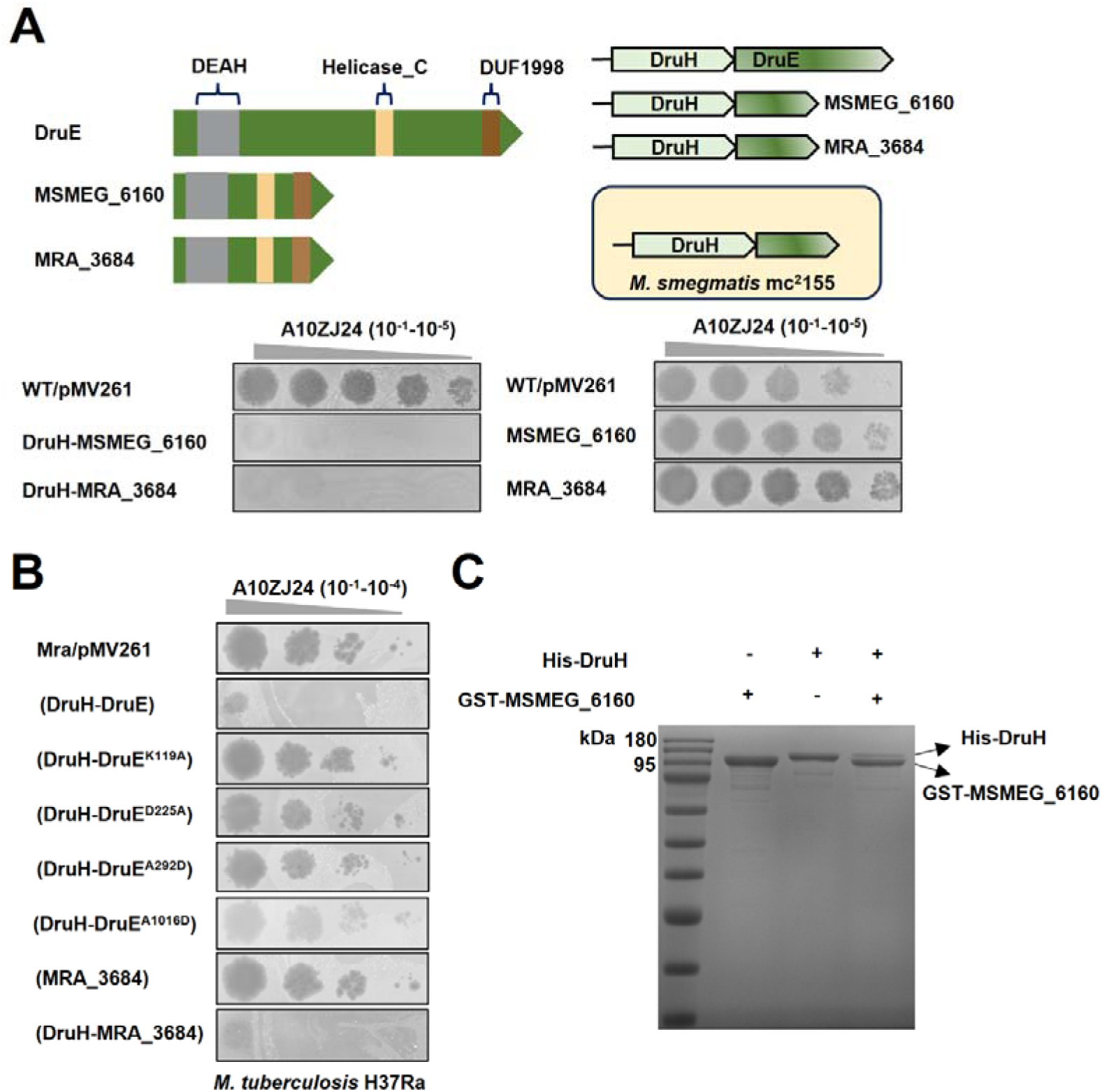
Antiphage activity assays for two mycobacterial DruE homologs. (A) DruE homologous protein and its anti-phage activity. The *M. smegmatis* MSMEG_6160 protein and the *M. tuberculosis* MRA_3684 protein have highly conserved domains with DruE (upper left). DruE in the type III Druantia system containing the native promoter was replaced with MSMEG_6160 or MRA_3684 and then cloned into *M. smegmatis* (upper right), 10-fold serial dilutions of plaque assays with phage A10ZJ24. WT/pMV261 and DruH-MSMEG_6160, DruH-MRA_3684, MSMEG_6160, MRA_3684represent wild-type strain and corresponding overexpression strains, respectively. (B) Plaque formation ability assays for phage A10ZJ24 on the lawns of different *M. tuberculosis* strains. Mra/pMV261 and (DruH-DruE) represent the wildtype strain and the type III Druantia-expressing *M. tuberculosis* strains, respectively. (MRA_3684), and DruH-MRA_3684 represent the wild type strain, MRA_3684-overexpression and DruH-MRA_3684 co-expression strain, respectively. The others represent overexpressing strains of the type III Druantia system with a site-specific mutation of the conserved amino acid residues of DruE, respectively. (C) GST pull-down assays for the interaction between DruH and MSMEG_6160. See also Figure S5 and S6.

To further determine whether the heterologous expression of type III Druantia has similar antiviral function, the DruH-DruE gene cassette was transformed into *M. tuberculosis* and the transformed strain was attacked with phage A10ZJ24. Compared with the control strain containing only the vector, the DruH-DruE recombinant strain showed significant resistance to the phage. In contrast, the DruH-DruE strains containing mutations in key amino acid sites of DruE (K119A, D225A, A292D, and A1016D) lost this resistance (Figure 3B). Strikingly, when DruH-MRA_3684 was transformed into *M. tuberculosis* to test its antiviral activity, the recombinant strain also showed significant resistance to the A10ZJ24 phage (Figure 3B).

Given that DruE is challenging to express and purify, and MSMEG_6160 can functionally substitute for DruE in conferring phage resistance, we next sought to determine whether DruH physically interacts with MSMEG_6160 using pull-down assays. Our results demonstrate that DruH specifically binds to GST-MSMEG_6160 but not to the GST tag alone (Figure 3C and Figure S5F). Furthermore, mass spectrometry analysis confirmed the presence of peptide fragments corresponding to both DruH and GST-MSMEG_6160 (Figure S6A and S6B), providing an evidence for the physical interaction between DruH and MSMEG_6160.

Therefore, these results indicate that two DruE homologs, *M. smegmatis* MSMEG_6160 and *M. tuberculosis* MRA_3684, can replace DruE from the DruH-DruE gene cassette to exert antiphage function. These recombinant hybrid systems can exhibit good antiviral function in different host bacteria such as *M. smegmatis* and *M. tuberculosis*.

### Type III Druantia system prevents phage DNA replication

Host bacteria can usually fight against phage infection by inhibiting adsorption, injection, replication, or activating abortive infection. In order to explore the defense mechanism of the type III Druantia system, we first compared the differences in phage adsorption rates between wild type and DruH-DruE strains. As shown in Figure 4A, no significant difference was observed for the adsorption rate between these two strains after measuring the phage adsorption at 30, 60, and 90 minutes after phage infection. This indicates that type III Druantia does not resist phage infection by blocking adsorption. Subsequently, we measured the changes in the number of intracellular phage genomic DNA in *mycobacterium* at different infection time points to observe the effect of DruH-DruE on phage DNA replication. As shown in Figure 4B, characterized by the essential genes of bacteriophages, the relative abundance value of phage DNA in DruH-DruE strain was only about 1.5 at 60 minutes after infection. In comparison, the wild type strain had the highest phage DNA abundance value of 13.5, which was nearly 9 times higher than that of DruH-DruE strain (Figure 4B), indicating that the phage DNA replication in DruH-DruE strains was prevented.

**Figure 4.**
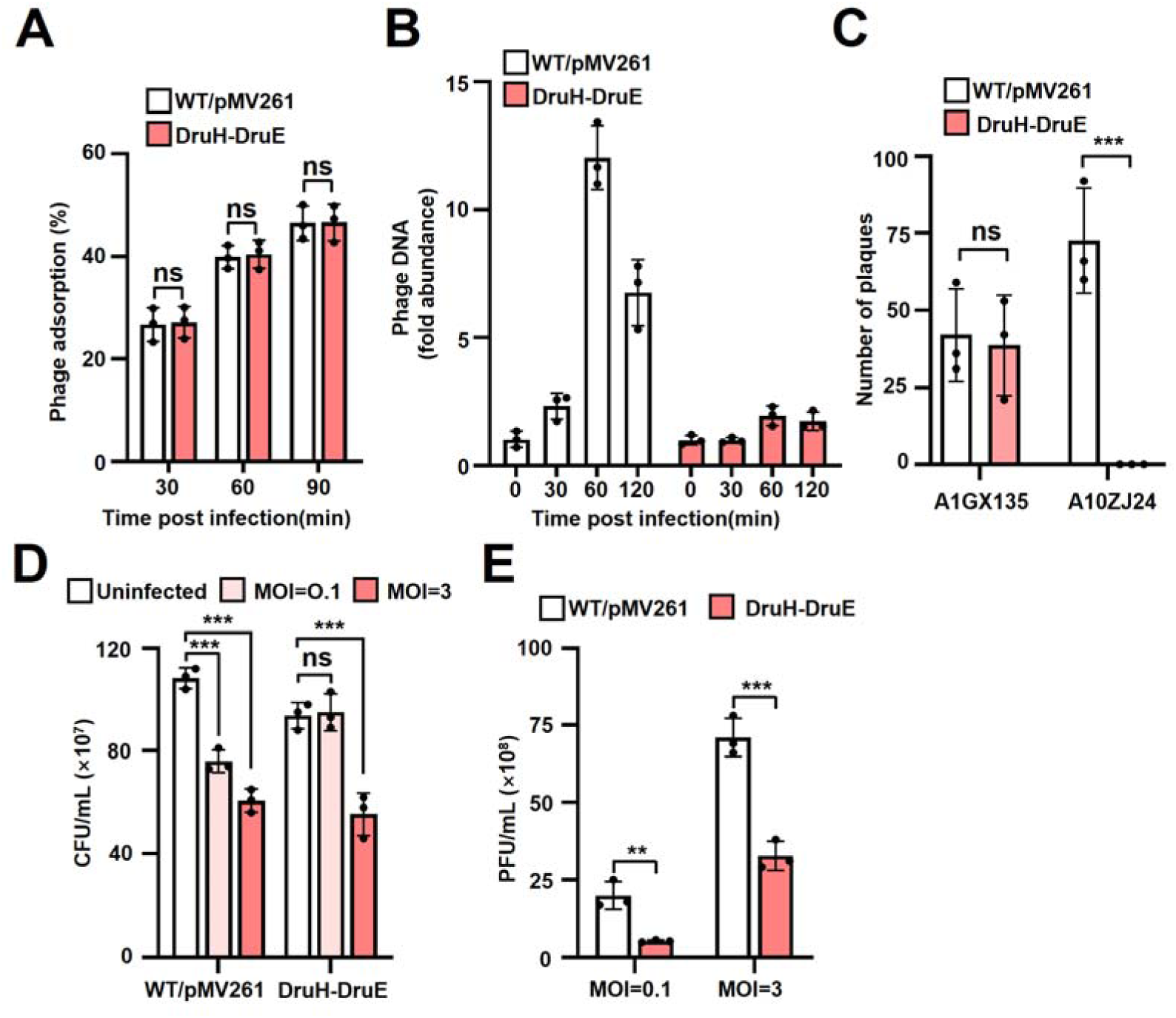
Type III Druantia prevents phage replication and activates abortive infection. (A) Phage adsorption assays for A10ZJ24 on the *M. smegmatis*. No significant difference was observed for the adsorption rates between type III Druantia-containing and the wild type *M. smegmatis* strains at 30, 60, and 90 minutes after infection. Error bars represent means ± SD. ns, no significant difference; unpaired two-tailed Student’s *t* test. (B) qPCR assays for relative abundance of phage DNA at various time points following phage infection. For all experiments, the mean ± SD of triplicate measurements are shown as a representative of at least three independent trials. (C) DNA electroporation assays. The extracted genome of phage A10ZJ24 was transferred into the type III Druantia overexpression strain by electroporation. A phage A1GX135 that can infect the DruH-DruE strain was used as a control, and the number of phage plaques was then assessed. Error bars represent means ± SD. ns, no significant difference; \*\*\**p*□<□0.001; unpaired two-tailed Student’s *t* test. (D) Assays for CFU/mL recovered from uninfected and different MOI (MOI=0.1, MOI=3) phage-infected cultures. WT/pMV261 and DruH-DruE represent the wildtype strain, and the type III Druantia system overexpression strain, respectively. Error bars represent means ± SD. \*\*\**p*□<□0.001; unpaired two-tailed Student’s *t* test. (E) Assays for plaques arising from phage (MOI=0.1 or MOI=3) infected cultures plated on the wild type lawn after removal of free phages in the medium. The DruH-DruE strain produced significantly lower PFU/mL than the control bacteria, regardless of exposure to high or low multiplicity of infection. PFU/mL values are the mean of three biological replicates; error bars represent means ± SD. \*\**p*□<□0.001; \*\*\**p*□<□0.001; unpaired two-tailed Student’s *t* test.

To further determine whether phages have replication defects due to the inability to inject genomic DNA into mycobacteria, we next conducted electroporation experiments by artificially injecting phages’ genomic DNA into bacterial cells to observe whether phages can successfully complete their subsequent life cycle. As shown in Figure 4C, the control phage A1GX135 is insensitive to the defense system. When its genomic DNA is electroporated into the DruH-DruE strain, no significant difference was observed in the number of plaques produced compared to the wild type strain, indicating that the electroporation experimental system worked well (Figure 4C). However, under similar conditions, no plaque formation was observed when the genomic DNA of the sensitive phage A10ZJ24 to the defense system was electroporated into the DruH-DruE strain. This indicates that the abnormal DNA replication in DruH-DruE strains infected with phage A10ZJ24 is not caused by DNA injection defects, but is most likely due to the genome being cleared by some mechanism after entering the DruH-DruE strain.

These results indicate that type III Druantia can prevent the replication of phage DNA during the infection.

### Type III Druantia system triggers phage abortive infection in ***M. smegmatis***

Given that the expression of the DruH gene alone affects the normal growth and even death of *M. smegmatis* strains (Figure S4A), we speculate whether type III Druantia may also prevent successful phage infection by activating abortive infection. We then tested this hypothesis. Abortion infection is a process involving premature death or growth arrest of infected cells, which prevents phage replication and spread to nearby cells (34). Therefore, when susceptible wild type cells are infected, it can lead to a decrease in CFU/mL. As shown in Figure 4D, when infected in liquid medium, exposure to high multiplicity of infection (MOI=3.0) resulted in a significant decrease in CFU/mL values from uninfected to infected cultures, independent of the strain (Figure 4D), indicating that the infected DruH-DruE mycobacterial cells cannot survive after phage infection. However, when exposed to low multiplicity of infection (MOI=0.1), the CFU/mL values of wild type cultures were significantly reduced from uninfected to infected cultures, while DruH-DruE strains were able to survive, with no significant difference in CFU/mL values compared to uninfected DruH-DruE cells (Figure 4D). Furthermore, we evaluated whether infected DruH-DruE cells released live phage offsprings. The results showed that the DruH-DruE strain produced significantly lower PFU/mL than the control bacteria, regardless of exposure to high or low multiplicity of infection (Figure 4E), indicating that the defense system prevents the release of phage offspring during phage infection, thus making it a typical phenotype of abortive infection.

Based on these results, the Type III Druantia system defends against phage infection by preventing phage DNA replication and inducing abortive infection.

### Type III Druantia system prevents phage DNA cyclization

Shortly after the injection, phage genomic DNA undergoes cyclization, which is essential for phage lysis or replication (35). Therefore, we used PCR amplification method to detect whether the type III Druantia system affects phage cyclization. Firstly, we confirmed that the phage A10ZJ24 did not produce lysogens, and DruH-DruE strain does not affect the phage entry into a lysogenic state. As shown in Figure 5, even after 2 hours of infection with phage A10ZJ24, no specific hybrid DNA fragments were amplified from either the wild type strain or the DruH-DruE strain after lysing (Figure 5A and 5B upper panel). However, we subsequently found that the DruH-DruE strain significantly affects the phage DNA cyclization. As shown in Figure 5B, a 875 bp specific DNA fragment produced by phage cyclization can be amplified in the wild type strain, but this fragment cannot be amplified from the DruH-DruE strain (Figure 5B, lower panel and Figure S7). Therefore, the type III Druantia system prevents phage DNA cyclization and further affects phage DNA replication.

**Figure 5.**
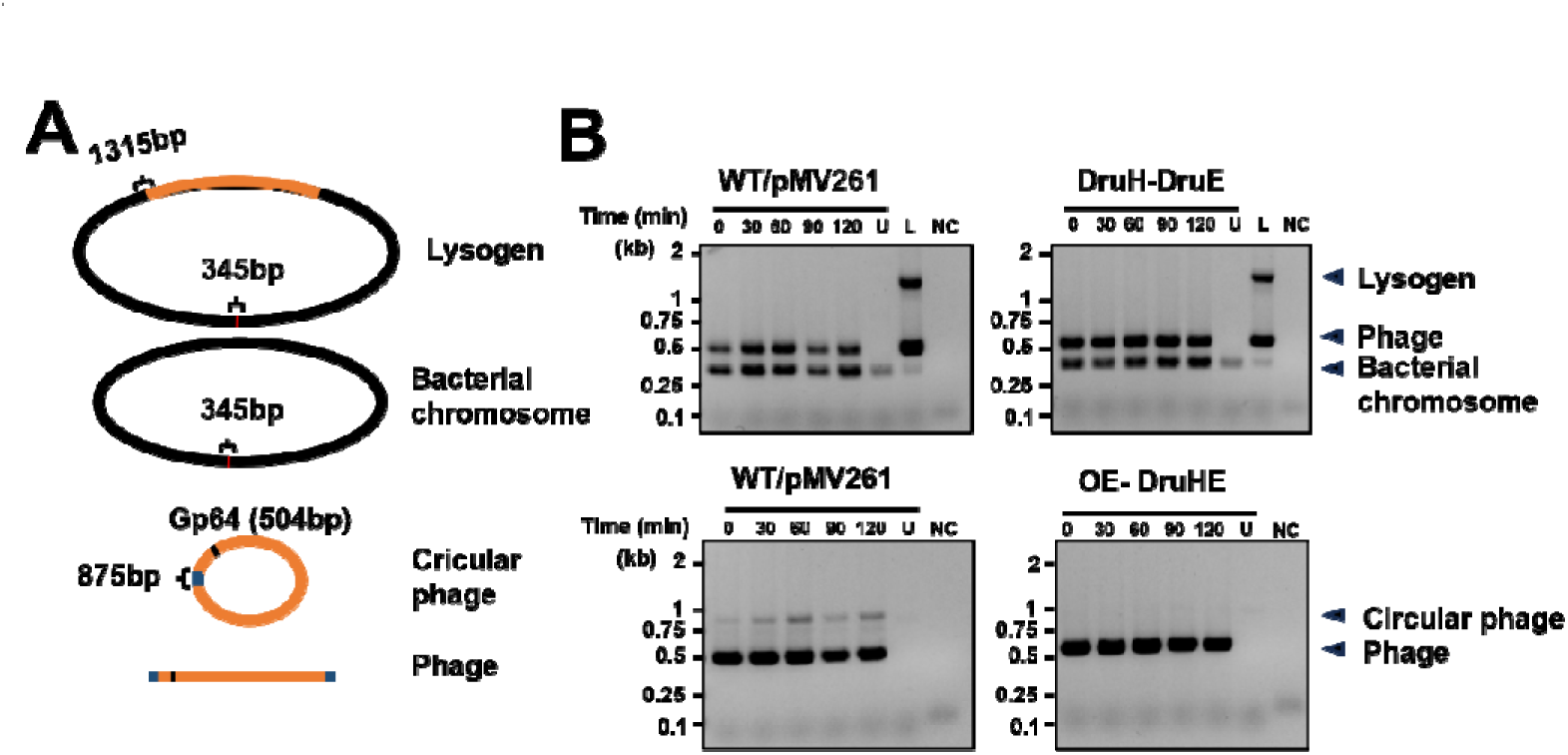
DruH-DruE prevents phage DNA cyclization. (A) Schematic representation of amplified DNA fragments in the experiments. Based on the sequencing results of lysogenic strains, specific primers were designed to detect the lysogenic status of the phage. The upstream primers were positioned within the *Mycobacterium* genome, while the downstream primers were located inside the phage genome, enabling the amplification of a 1315 bp fragment. Furthermore, the phage A10ZJ24 was identified using the *gp64* gene, which yielded a 504 bp product. To assess phage circularization, primers were designed to flank the Overhang Sequence (CGGCCGGGTAA), resulting in the specific amplification of an 875 bp fragment. Amplicons for the bacterial DNA is 345bp. (B) The type III Druantia system inhibits phage Genome circularization. WT/pMV261 and DruH-DruE represent the wildtype strain, and the type III Druantia overexpression strain respectively. Agarose gel of multiplex PCR with 2 or 3 primer sets, aimed to detect bacterial DNA, phage DNA and lysogen. Lanes are marked with minutes post infection; U lane is the uninfected control, L lane is the lysogen, NC lane is the blank control. Primers designed on either side of the phage cos site for detection of phage circularization. See also Figure S7

### Phage-encoded Gp25 activates type III Druantia system

Since the expression of type III Druantia system is not toxic to host bacteria in the absence of phage infection, we infer that this defense system may be activated by phages. To test this issue, the phage escapees against type III Druantia system were repeatedly enriched (Figure 6A). Fortunately, after multiple rounds of screening, a mutant phage Mut-A10ZJ24 was successfully isolated, as shown in Figure 6B. The phage can effectively infect DruHE-expressing strains, and its plaque formation capacity on double-layer plates is very similar to that on wild type strains (Figure 6C). Further sequencing analysis showed that Mut-A10ZJ24 had mutations within three genes, including *gp23* (encoding a tape measure protein), *gp24* (encoding a minor tail protein) and *gp25* (encoding a minor tail protein) (Figure 6D and Figure S8A). To further identify the type III Druantia system activators among these candidate genes, we co-expressed each of these three genes together with DruH-DruE to examine if the DruH-DruE toxicity can be activated. The results showed that the expression of wild type *gp25* resulted in significant toxicity of co-expressed DruH-DruE to host mycobacteria. By contrast, the other two genes, *gp23* and *gp24*, had no effect on bacterial growth under the same conditions (Figure S8B and S8C). Furthermore, no effect on bacterial growth was observed (Figure 6E) when introducing mutations into the DruH-DruE gene cassette and co-expressing with *gp25*, or co-expressing mutant *gp25* with wild type DruH-DruE gene cassette.

**Figure 6.**
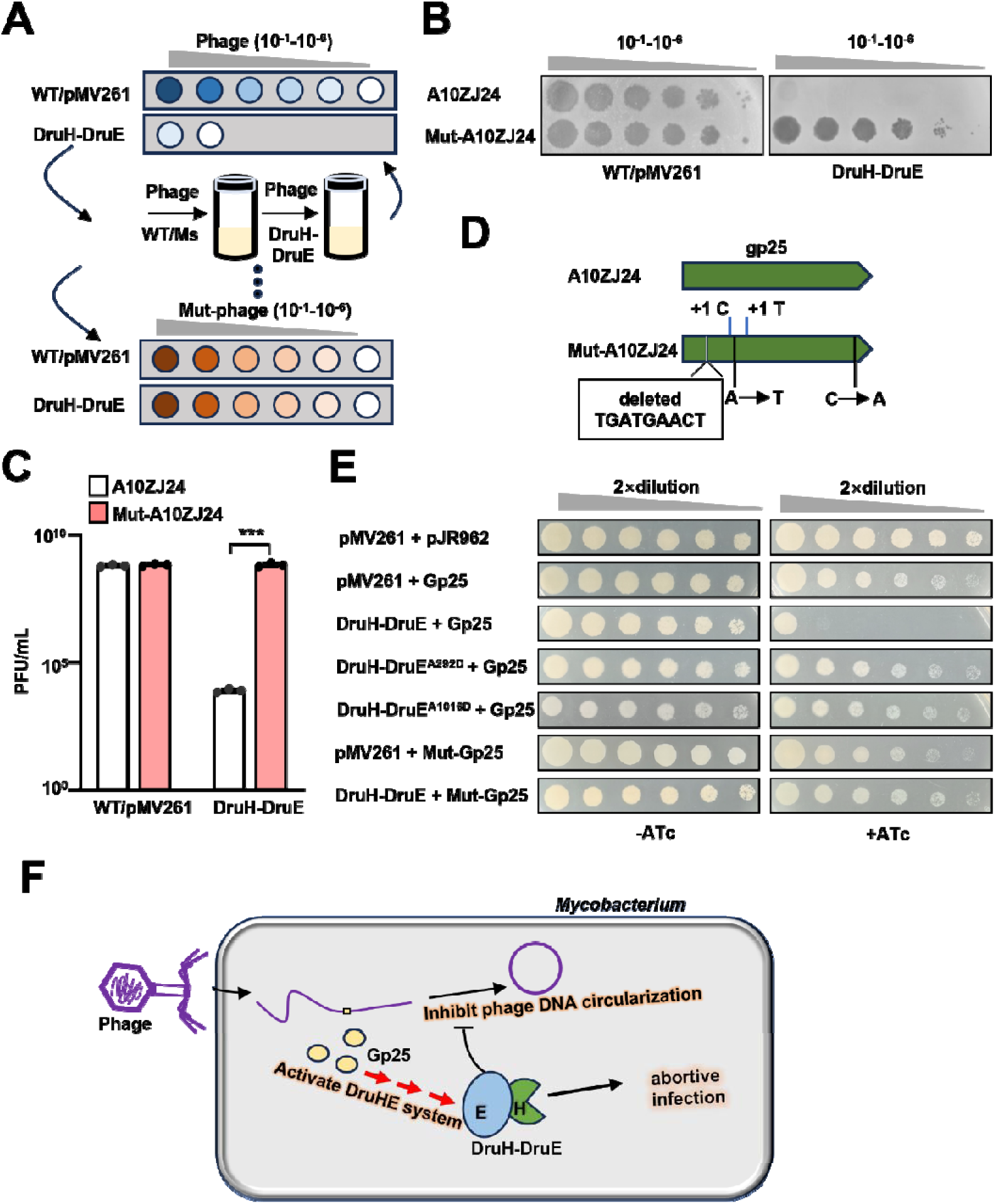
Type III Druantia system is activated by the minor tail protein. (A) Schematic representation of phage evolution experiments. Individual phage plaque formed on the DruH-DruE strain is collected for screening phage mutants. The collected plaques are tested for their ability to overcome the defense system using serial dilution plaque assays. (B) 10-fold serial dilution assays for the plaque formation ability of phage A10ZJ24 and its mutants. (C) Evaluation of phage plaque formation efficiency of mutant phage (Mut-A10ZJ24). The number of phages per 1 mL (plaque-forming unit [PFU]/mL) was assessed after adding phage suspension to the double-layer agar plates at 24 h. WT/pMV261 and DruH-DruE represent the wild-type strain, and the type III Druantia overexpression strain respectively. Error bars represent means ± SD. \*\*\**p*□<□0.001; unpaired two-tailed Student’s *t* test. (D) Schematic diagram of the mutation in the minor tail protein *gp25* gene of phage A10ZJ24, with mutations in the escape clones from B labelled. (E) Expression of a phage-encoded Gp25 activates toxicity in a strain that contains the Type III Druantia system. Serial dilutions of cells were incubated in conditions with (+) or without (−) anhydrous tetracycline, bacterial viability was measured for strains in which the type III Druantia system was co-expressed with a phage gene (*gp25*) or co-expressed with a phage mutation gene (Mut-*gp25*). pMV261-pJR962 represents the wild-type strain. pMV261 + Gp25 represents the ATc-induced strain for Gp25 expression strain with an empty pMV261 vector. pMV261 + Mut-Gp25 represents the the ATc-induced strain for mutant phage gene (Mut-gp25) expression. DruH-DruE + Gp25 represents the DruH-DruE strain with recombinant vector pJR962-*gp25*, in which Gp25 expression is induced by ATc. pMV261 + Mut-Gp25 represents the ATc-induced strain for mutant phage gene (Mut-*gp25*) expression strain with an empty pMV261 vector. DruH-DruE^A292D^ + Gp25 and DruH-DruE^A1016D^ + Gp25 represent the DruH-DruE mutant strains with recombinant vector pJR962-*gp25*, in which Gp25 expression is induced by ATc. (F) Model of antiphage defense by the type III Druantia system. After phage genome injection, the production of small tail protein activates the type III Druantia system, a defense system that prevents phage circularization and inhibits phage replicationandleading to abortive infection. See also Figure S8.

Therefore, these results indicate that the phage-encoded Gp25 can activate the type III Druantia defense system to co-induce lethality of host mycobacterial cells, which is consistent with the previous observation that phage infection promotes the occurrence of abortive infection in DruH-DruE strains.

## Discussion

The antiphage mechanism of type III Druantia system remains unclear. In this study, we provided evidence to show that the type III Druantia is a two-component antiphage defense system. Both DruH and DruE are essential for fighting against phage infection and the defense function also depends on the interaction between DruH and DruE. Our findings support an antiviral defense model for the type III Druantia system (Figure 6F), in which the defense system is activated by the small tail protein of phages during infection. Thereafter, the DruH prevents phage genomic DNA cyclization, which halts phage DNA replication and finally triggers abortive infection for defending against broad-spectrum phages. Our study expanded our understanding of the immune mechanisms in prokaryotic cells.

In this study, we noticed the high similarity between the type III Druantia and Hachiman systems, which are both composed of dual genes and contain a hypothetical protein and helicase protein. However, the obvious difference is that DruH encodes four times more amino acids than HamA, and the helicase DruE (1667aa) and HamB (1174aa) differ by nearly 500 amino acids. Although the composition genes of these two defense systems differ greatly in size, there seems to be significant similarity in their antiphage mechanisms, both of which can cause Abi reactions after phage infection (Figure 3D and 3E), which may suggest inherent connections and conservation between different defense systems.

The DruE gene of type III Druantia system encodes a larger protein with 1667aa containing three conserved motifs (Figure S1A). According to our site-directed mutagenesis experiments, the DruE helicase domain, DUF1998, and DExD/H box motif are necessary for its antibacterial activity (Figure 2B). Recent studies have described DruE helicase as a potential unique branch of the SF2 helicase family (6). In addition to the restrictive modification system, the phage defense module also contains various SF2 helicases, and the DExD/H box helicase domain is also related to the previously reported prokaryotic Argonaute system (36–38). In addition, the functioning of the Type I CRISPR Cas system requires the SF2 helicase/nuclease protein Cas3 (39, 40), while the Type III and V BREX systems include a helicase BrxHII with a DEXD-RapA domain (7), and the helicase/nuclease Hna protein N-terminus contains the superfamily II (SF2) motif (21). It is interesting that type III Druantia, like Hna, can cause Abi response in infected cells. But the phage resistance provided by the DruE protein alone is significantly weaker than that of the intact Druantia system (Figure S4B). Although the specific function of the DruE gene remains to be further understood, it is interesting to note that we have discovered a homologous protein SftH encoded by MSMEG_6160 in the genome of *M. smegmatis*, which shares the same conserved domain between two proteins (Figure 3A and Figure S5A, S5B). SftH has been reported as a monomeric DNA dependent ATPase/dATPase that can translocate 3’ to 5’ on single-stranded DNA and exhibits 3’to 5’ helicase activity (24). In this study, through gene replacement, we confirmed that SftH can replace the DruE gene to form a phage defense system similar to the DruHE system with DruH, which suggests that DruE may have a considerable degree of functional homology with SftH protein.

Another interesting finding from this study is that the small tail protein encoded by phages can activate the function of the type III Druantia defense system. One of the main strategies for phages to escape the host defense system is through mutations in their structural proteins (41). Previous studies have shown that mutations in capsid proteins enable phages to overcome multiple defense systems. For example, CapRelSJ46 is activated by the main capsid protein of SEC Φ 27 (42). In addition, DSR2 protein can directly recognize phage tail tube proteins and activate their NADase activity, thereby consuming intracellular NAD^+^ and causing Abi reaction (43). In the present study, our results indicate that phages can also evade the defense of type III Druantia system through mutations in their small tail protein gene, and even expressing small tail protein alone in cells expressing the defense system is sufficient to activate Druantia’s defense response, causing the death of host bacterial cells (Figure 6E and Figure S8B, S8C). Our research has to some extent enriched the ways phage immune escape occurs, but its exact activation mechanism still needs further clarification.

In summary, this study identified for the first time the antiviral mechanism of the type III Druantia system, greatly enriching the library of mycobacterial defense systems against phages and expanding our understanding of the molecular mechanisms of the prokaryotic immune system.

## Materials and Methods

### Bacterial strains and media

Liquid cultures of *M. smegmatis* mc^2^155 were grown in Middlebrook 7H9 media at 37 °C at 160rpm or in 7H10 medium (BD Difco) containing 0.5% glycerol at 37°C. *E. coli* strains were grown in LB media at 37 °C at 160rpm. Whenever applicable, media was supplemented with chloramphenicol (34 μg/ mL), tetracycline hydrochloride (12.5 μg/mL) or kanamycin (30 μg/mL) to ensure plasmid maintenance. Mycobacteriophages were isolated from different soil samples in China, and the wild type *M. smegmatis* strain was used for their propagation.

### Construction of recombinant *M. smegmatis* strains

The pMV261 vector was used to construct recombinant plasmids containing different mycobacterial or phage genes following previous procedures (44). Briefly, type III Druantia system containing the native promoter region obtained by PCR, and was inserted into the pMV261 vector between the *Xba* I and *Hind* III, while other mutant amplicons were also separately assembled into the vector between the *Xba* I and *Hind* III restriction sites using the Uniclone One Step Seamless cloning kit (Genesand, China). The resulting plasmids were transformed into the *M. smegmatis* strains and plated on 7H10 medium supplemented with 30□μg/mL kanamycin. To construct Druantia and phage gene co-expression strain, the above-mentioned pMV261 vector and an ATc-inducible expression vector pLJR962 were used (31). Amplified phage gene were fused into the modified pLJ962 vector between the *Cla*I and *Not*I restriction sites to obtain the recombinant plasmid. Next, the two recombinant plasmids were co-transformed into *M. smegmatis* mc^2^155, and plated on 7H10 medium containing 30 μg/mL kanamycin and 50 μg/mL hygromycin.

### Isolation of phage mutant strains

Defense systems expressing in *M. smegmatis* provide protection against phage infection, as observed using serial dilution plaque assays. Phage plaques that are formed on bacteria expressing the defense system are collected to screen for escaping phage mutants. The experimental flow is shown in Figure 6A, the collected phage plaques were first co-incubated with wild type *M. smegmatis* overnight, followed by co-incubation of the filtrate with the defense system-containing strain overnight, and so on for several times until mutant phages sensitive to the defense system were obtained. The genome of the mutant strain was extracted and sequenced by BGI group.

### Phage adsorption assays

Phage adsorption assays were performed according to previous procedures (30, 45). *M. smegmatis* was cultured at 37°C until OD_600_ = 1.0 and then centrifuged. Cells were resuspended in 7H9 medium without Tween 80 and phage was added to 10 mL of culture at an MOI of 0.01. Subsequently, the culture was shaken slowly at 60 rpm at 37 °C, and 1 mL of the sample was removed and centrifuged at 30 min, 60 min, and 90 min post-infection. Subsequently, 100 μL of the supernatant was taken and measured by double layer plaque assay to determine reference phage concentration.

### Protein expression and purification

The gene encoding DruH protein was amplified by PCR using the specific primers listed in Supplementary Table 1. The amplicons were separately assembled into pRSF-Dut vectors and then transformed into *E. coli* BL21 (DE3). Cultures were grown in LB medium containing 30 μg/m kanamycin at 160 rpm at 37°C, until the OD_600_ reached about 0.8. Cultures were cooled on ice for 1 h, and then induced with 0.5 mM IPTG for 20 h at 16°C. All subsequent procedures were performed at 4°C and referred to the method described previously (24). The eluted proteins were dialysed in low-salt buffer (pH=7.5, 25 mM Tris-HCl, 100 mM NaCl, 1 mM DTT, 10% glycerol) for 3 hours and stored at −80°C. Protein concentration was determined using the Coomassie Brilliant Blue assay.

### qPCR assays

*M. smegmatis* mc^2^ 155 strains were cultured at 37°C until OD_600_ = 1.0 and then centrifuged. The cells were resuspended in 10 mL of 7H9 medium without Tween 80 and phage with an MOI of 1 was added. Subsequently, the cells were incubated at 37 °C for 30 min with slow shaking at 60 rpm, and centrifuged. Next, the cells were resuspended in 100 mL of 7H9 medium without Tween 80, and Immediately 10 mL aliquots were removed as the zero moment and stored at −80 °C, and 10 mL of the sample was centrifuged at 30 min, 60 min, and 120 min post-infection, and their total DNA was extracted. DNA extraction was conducted as described previously (46). Each qPCR reaction (25 μL) contained 500 ng of total DNA as template, 0.4 nM of phage-specific primers or host-specific primers (Supplementary Table 1) and 2X SYBR Green Mix (keepbio). The qPCR reaction procedure was as described previously (5). Phage DNA copy number was normalized against host values using the *16S rRNA* gene. Relative DNA abundance was determined using the 2^−ΔΔCT^ method.

### Bacterial two-hybrid assay

BacterioMatch II Two-Hybrid System Library Construction Kit (Stratagene) was used to detect protein–protein interactions between DruH and DruE as described previously (47). Briefly, positively growing co-transformed strains were screened on selective screening medium plates containing 5 mM 3-amino-1, 2, 4-triazole (3-AT) (Stratagene), 8 μg/mL streptomycin, 15 μg/mL tetracycline, 34 μg/mL chloramphenicol and 30 μg/mL kanamycin. Co-transformed strains containing pBT-*LGF2* and pTRG-*Gal11P* (Stratagene) were used as a positive control (CK^+^) to test the expected growth on the screening medium. Co-transformed strains containing the empty vectors pBT and pTRG were used as the negative control (CK^−^).

### DNA electroporation assay

The genome DNA of phages was extracted using the phenol-chloroform method as described earlier (48). The experimental operations are referenced as previously described with additional modifications (49). Briefly, a total of 200 ng of phage genomic DNA was electroporated into WT/PMV261 and DruH-DruE strains, respectively, and supplemented with 900□μL 7H9 medium, followed by shaking at 160□rpm at 37°C for 1□h, and then collected. The cells were re-suspended with 200□μL of 7H9 medium. Cells were added to 250 μL of wild-type *M. smegmatis* (WT/pmv261) and then mixed with 5 mL of LB medium supplemented with 0.6% agar. The mixture was plated on 7H10 medium and incubated at 37°C for 24 h, the plaque number was counted.

### Phage spotting assays and PFU measurements

The experiments were performed as previously described with additional modifications (42). To titre phage, dilutions of phage stocks were mixed with *M. smegmatis* and melted LB + 0.6% agar and spread on 7H10 medium plates and incubated at 37□°C for 24 h. For phage spotting assays, 1mL of a bacterial strain of interest (OD_600_□=□1.0) was mixed with 5 mL LB + 0.6% agar and spread on 7H10 medium plate. Phage stocks were then serially diluted in MP buffer (50□mM Tris-HCl pH 7.5, 150□mM NaCl, 10□mM MgSO_4_, 2 mM CaCl_2_), and 2□μL of each dilution was spotted on the bacterial lawn. Plates were then incubated at 37□°C for 24 h. PFU measurements were performed. To estimate free phage particle production from a single round of infection. Briefly, 100□μL of the diluted phages were mixed with 1□mL cultures of mycobacterial strain and 5□mL of LB + 0.6% agar. The mixture was then plated on 7H10 medium at 37°C for 24 h incubation, the number of plaques was counted. Data reported are the mean and individual data points from 3 biological replicates.

### Phage circularization assays

Genomic DNA sequencing of a lysogen containing phage A10ZJ24 was performed using Illumina sequencing to determine the DNA sequence of the A10ZJ24 phage, and the site of phage integration in the genome. The A10ZJ24 phage was determined to be integrated at a GAAGAG site within the *MSMEG_1635* gene of the *M. smegmatis* mc^2^ 155 bacterial genome. The experiments were performed as previously described with additional modifications (13). *M. smegmatis* mc^2^ 155 strains were cultured in 100 mL 7H9 medium at 37°C until OD_600_ = 1.0 and then centrifuged. The cells were resuspended in 50 mL of 7H9 medium without Tween 80 and phage with an MOI of 1 was added. 5 mL samples were taken immediately after infection (t=0) and at 30, 60, 90, 120 minutes after infection. During the infection, the culture was incubated with shaking slowly at 60 rpm at 37 °C. An uninfected control sample was taken before addition of phage. Samples were then centrifuged and the pellet was washed 3 times in PBS buffer to remove unabsorbed phages. Total DNA was extracted using Qiagen DNeasy Blood & Tissue kit (Qiagen 69504). Detection of phage lysogeny was performed using multiplex PCR as previously described by Goldfarb *et al* (7). Phage genome and bacterial genome were detected using their specific PCR primers (Supplementary Table 1), respectively. The lysogeny junction was detected using specific primer pair of Lysogen-F/Lysogen-F. Detection of phage circularization was done using primer pair of Circular-F/Circular-R (Supplementary Table 1).

### Monomer and multimer protein structure prediction by Alphafold

AlphaFold (32) is used to model the monomer structure, and PyMOL is employed to export and compute their average pLDDT scores. AlphaFold-Multimer (33) was employed for protein-protein interaction predictions using default parameters and a maximum template date of 2022-12-31 (--max_template_date=2022-12-31). PyMOL is employed to visualize the interaction interface.

### GST pull-down assay

The GST and GST-tagged MSMEG_6160 (GST-MSMEG_6160) proteins were expressed in *E. coli* BL21 (DE3) and purified. Following purification, the eluates were dialyzed overnight at 4 °C. For the pull-down assay, approximately 100 µg of GST or GST-MSMEG_6160 was immobilized onto 50 µL of glutathione-agarose resin, equilibrated, and incubated at 4 °C for 1 hour with gentle agitation, followed by three washes with PBST. Subsequently, approximately 300 µg of His-DruH was added to the immobilized GST-MSMEG_6160 or GST. The protein complexes were incubated at 4 °C under gentle rotation for 4 hours. After washing three to four times with PBST, bound proteins were eluted using elution buffer (10 mM glutathione in PBS, pH 8.0) and analyzed by SDS-PAGE.

### NanoLC-MS analysis

The proteins from the shifted band corresponding to the GST-MSMEG_6160 and His-DruH in the gel were extracted and processed according to previously published procedures (50). The samples were digested with trypsin and subsequently vacuum-concentrated to remove residual liquid. The resulting supernatant was loaded into an injection vial for analysis using an EASY-nLC 1000 system coupled to an LTQ-Orbitrap Elite mass spectrometer (Thermo Fisher Scientific, USA). The mobile phases consisted of 0.1% formic acid in aqueous solutions with varying acetonitrile concentrations: mobile phase A contained 2% acetonitrile, while mobile phase B contained 98% acetonitrile. A linear gradient elution program was implemented as follows: 2 - 12% B over 0 - 5 min, 12 - 20% B from 5 - 30 min, 20 - 32% B from 30 - 43 min, and 32 - 98% B from 43 - 58 min, maintained at a constant flow rate of 250 nL/min. Mass spectrometric analysis was performed with a scan range of *m/z* 350 - 1500. The Orbitrap analyzer achieved mass resolutions of 70,000 for full MS scans and 17,500 for MS/MS scans. Protein identification was conducted using Proteome Discoverer 2.5 software, with database searches specifically targeting peptide matches for DruH and MSMEG_6160 proteins based on the acquired NanoLC-MS spectra.

## Acknowledgements

This work was supported by the National Natural Science Foundation of China (32230002) and National Key R&D Program of China (2020YFA0907200).

