## Supplemental information for "Type III Druantia two-component antiphage defense depends on the DruH-DruE interaction for halting phage DNA cyclization and replication"

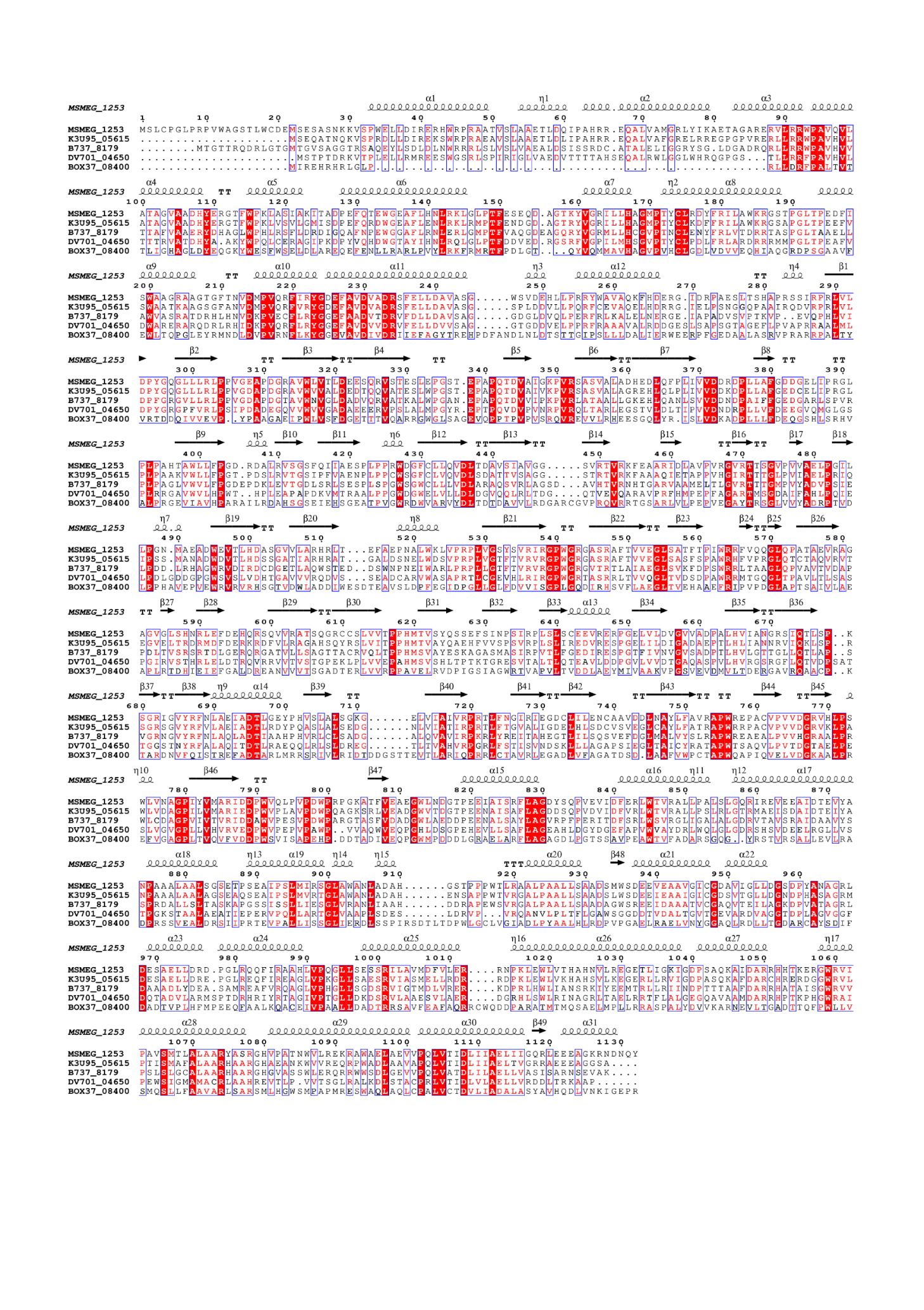

**Figure S1, related to Figure 1. Amino acid sequence conservation analysis of DruH in the Type III Druantia system.** The conservation of amino acids in DruH, a key component of the type III Druantia system, was analyzed and visualized using a color to represent the degree of conservation across homologous sequences.

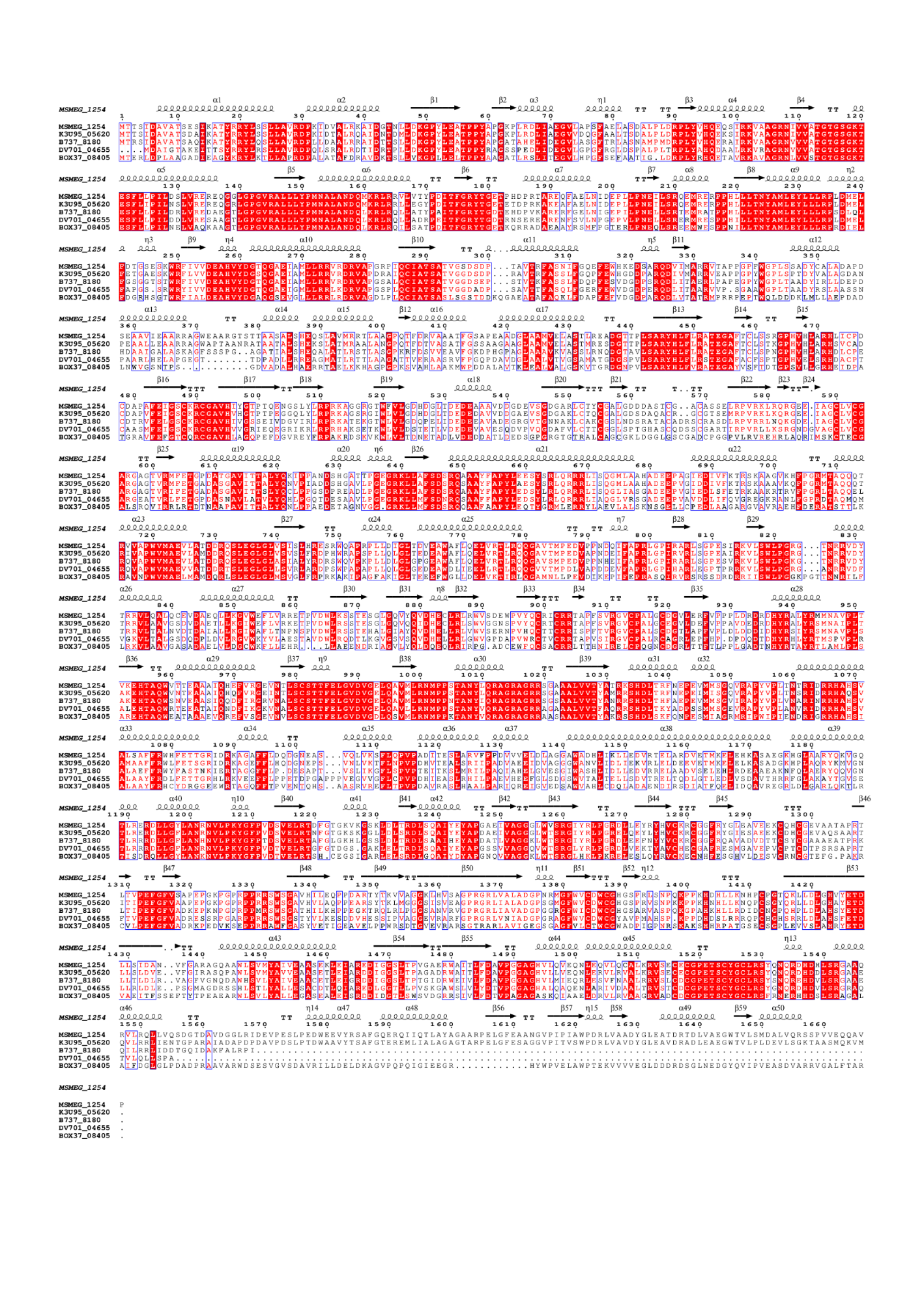

**Figure S2, related to Figure 1. Amino acid sequence conservation analysis of DruE in the Type III Druantia system.** The conservation of amino acids in DruE was analyzed and visualized using a color to represent the degree of conservation across homologous sequences.

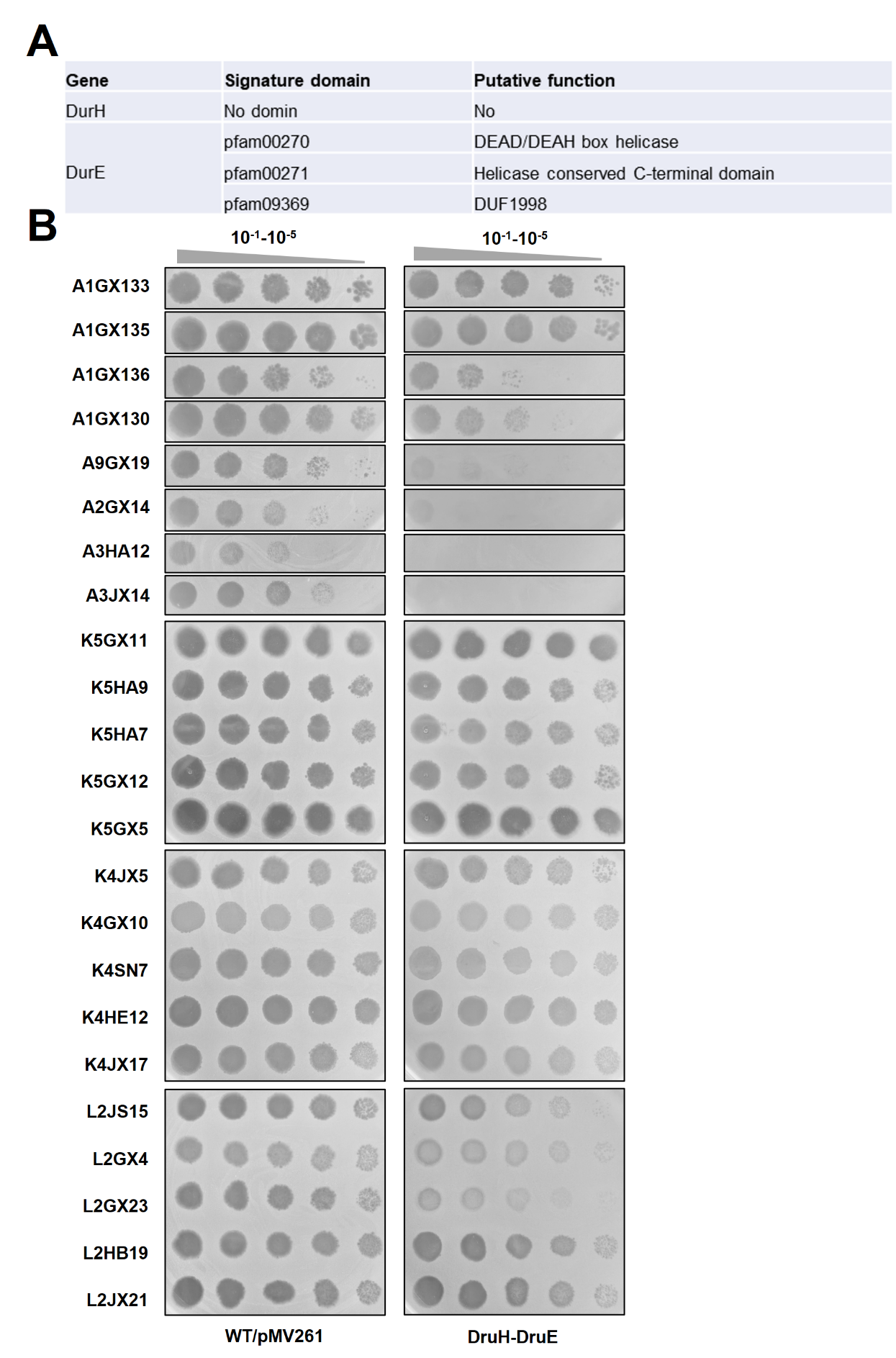

**Figure S3, related to Figure1. Phage phenotypes assays for the** **DruH-DruE overexpression strain.**

(A) Gene organization and domain annotation of the Type III Druantia system.The genetic architecture of the type III Druantia system, including its constituent genes and their associated functional domains, is depicted.

(B) Serial dilution assays for antiphage activity. Tenfold serial dilutions of a panel of genetically diverse mycobacteriophages were spotted onto lawns of *M. smegmatis* strains, including the wild-type (WT/pMV261) and the overexpression strain (DruH-DruE). The names of the tested phages are indicated on the left.
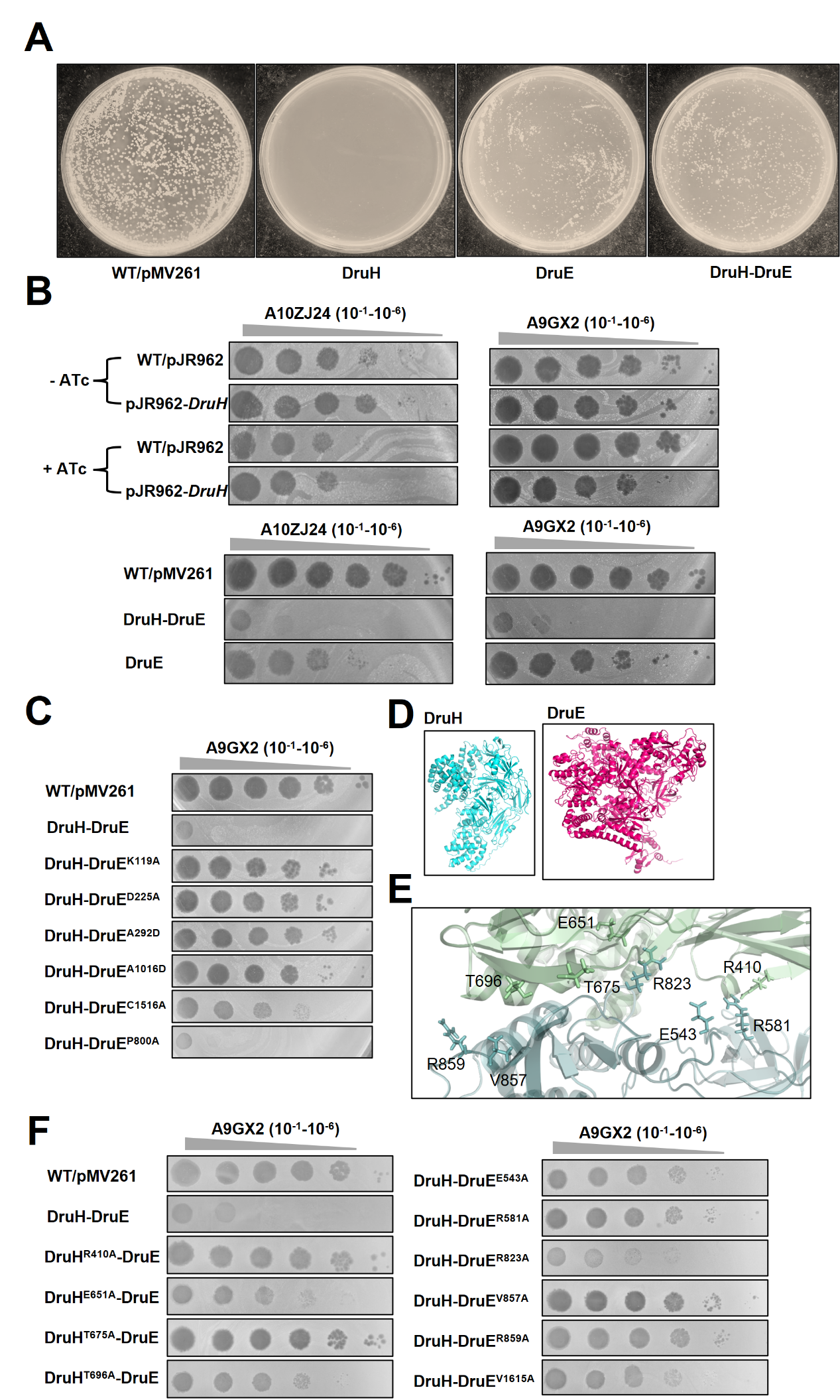

**Figure S4,** **related to Figure 2. Antiphage A9GX2 phenotype assays for type III Druantia system containing mutations in DruH or DruE.**

(A) Growth of different *M. smegmatis* strains on 7H10 solid medium. Strains included the wild-type (WT/pMV261) and DruH, DruE, and DruH-DruE overexpression strains.

(B) Antiphage activity of the type III Druantia system. Tenfold serial dilution plaque assays were performed to evaluate the infection efficiency of phages A10ZJ24 and A9GX2 on *M. smegmatis* strains expressing the type III Druantia system or individual components (DruH or DruE)..

(C) Functional analysis of DruE mutations in the Druantia system. The phage resistance of the type III Druantia system was tested using strains harboring mutations in DruE, revealing the contribution of specific residues to antiphage activity.

(D) Structural prediction of DruH and DruE. The AlphaFold-predicted structures of DruH and DruE were generated to elucidate their molecular architecture and potential interaction interfaces.

(E) DruH-DruE complex interface analysis. The AlphaFold-predicted structure of the DruH-DruE complex was analyzed to identify key residues at the interaction interface, providing insights into the molecular basis of their functional partnership.

(F) Functional impact of DruH-DruE interface mutations. Mutations at the DruH-DruE interface were introduced based on residues with protein-protein contacts. These mutants were tested to determine their effects on the system’s ability to confer phage resistance, highlighting the importance of specific interactions for antiphage activity.

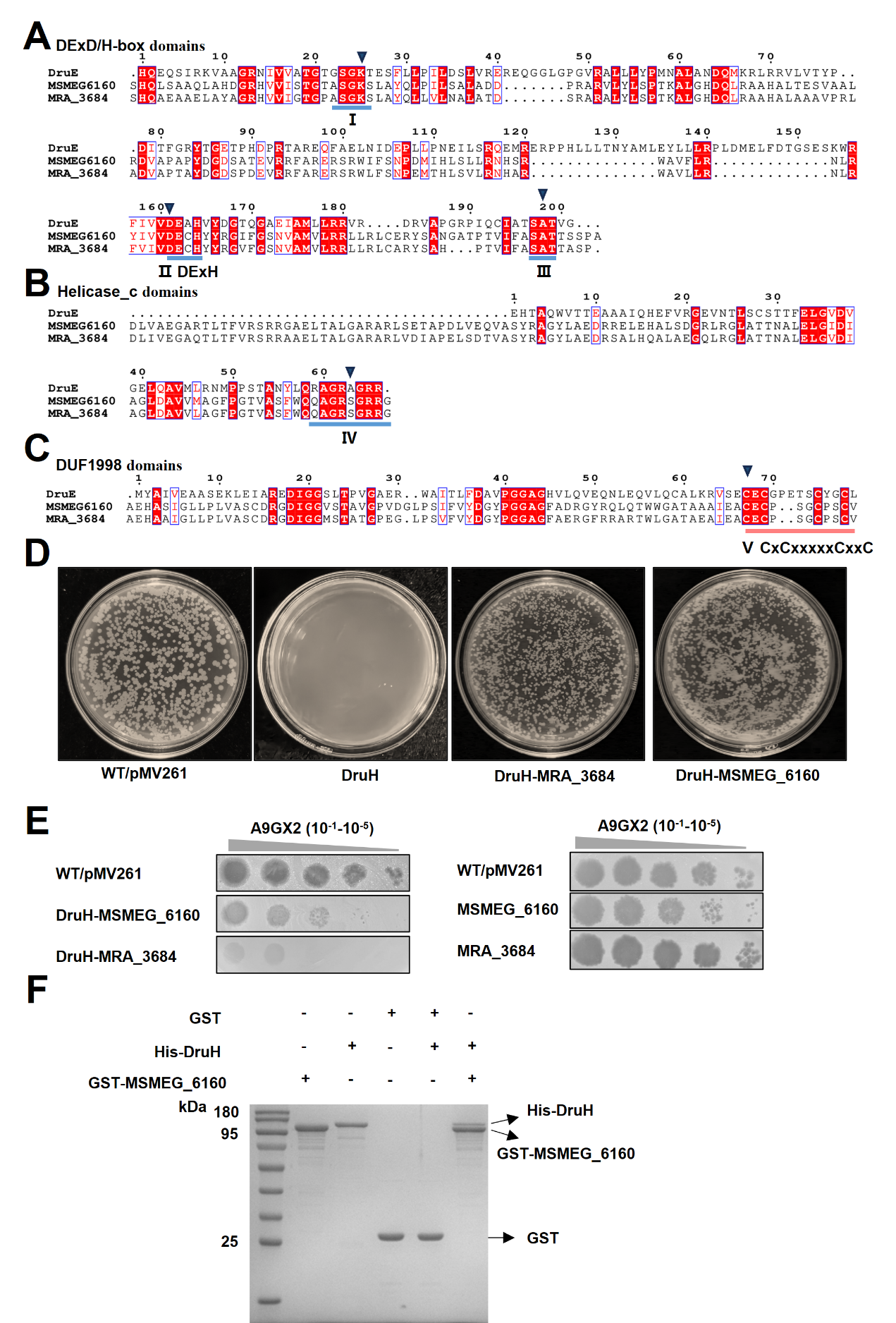

**Figure S5,** **related to Figure 3. Antiphage activity of DruE homologous protein in *Mycobacterium*.**

(A-C) Amino acid sequence conservation analysis of DExD/H-box, helicase_C and DUF1998 structural domains of DruE, MSMEG_6160, and MRA_3684Mra_3684. The superfamily II ATPase motifs I, II, III and Ⅳ are highlighted in blue, Ⅴ represents tetracysteine cluster. The triangles represent DruE point-mutation sites.

(D) Growth of different *M. smegmatis* strains on 7H10 solid medium. WT/pMV261 indicates the wild-type strain. WT/pMV261 represents the wild-type strain. DruH, DruH-MRA_3684Mra_3684, DruH-MSMEG_6160 represent *M. smegmatis* strains with the corresponding gene overexpression, respectively.

(E) Phage plaque formation ability assays. 2 µL of serial dilutions of A9GX2 phage were spotted on lawns of mycobacterial cells (OD_600_ =1.0) and plaque formation was assessed after 24h of incubation at 37 ℃.

(F) GST pull-down assays for the interaction between DruH and MSMEG_6160. The GST tag protein was used as a control.

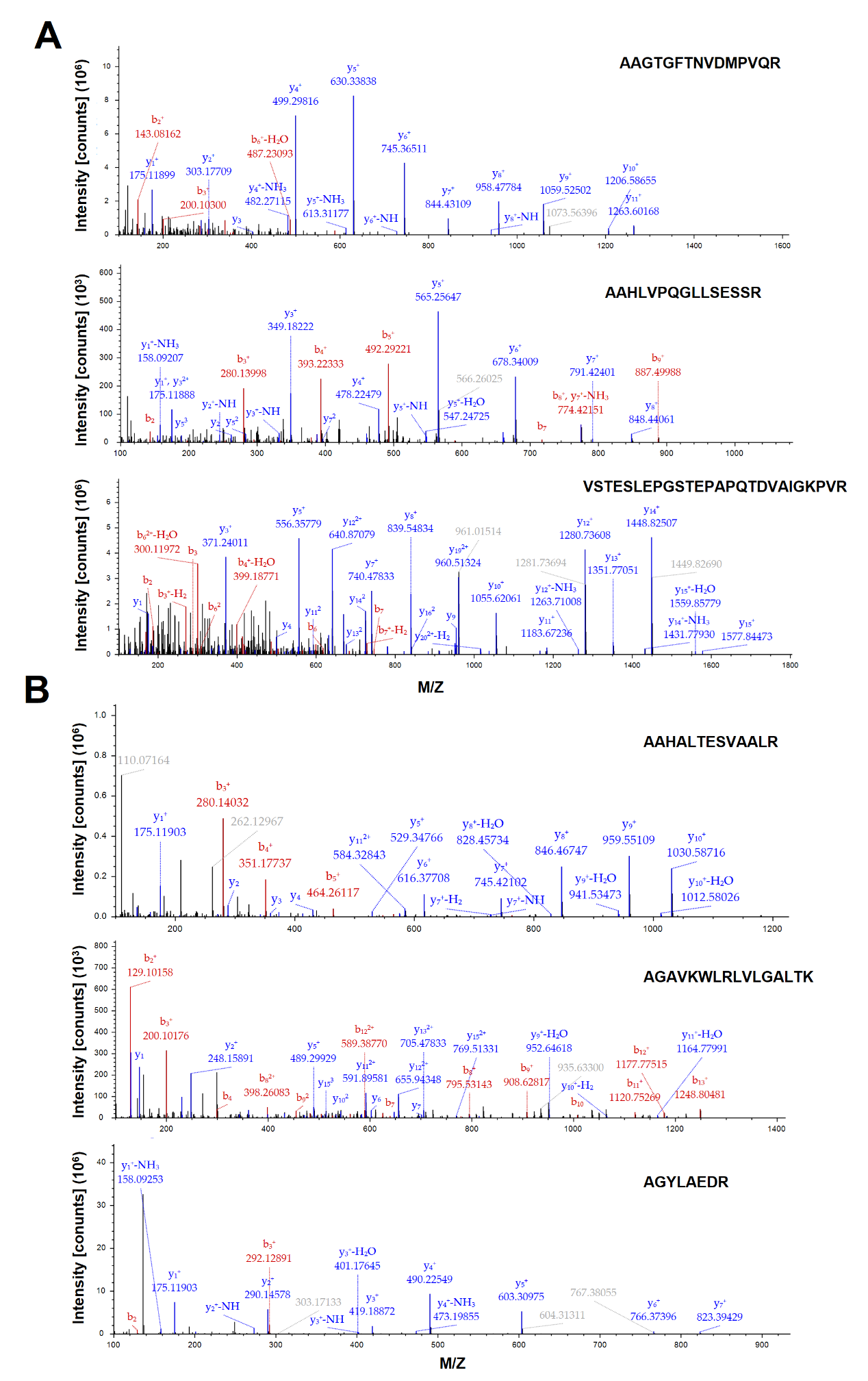

**Figure S6,** **related to Figure 3. Identification of pull-down protein samples by nano-liquid chromatography-mass spectrometry (NanoLC-MS).**

(A) NanoLC-MS assays for specific peptide fragments of DruH. DruH-specifc peptide fragments with the amino acid sequences AAGTGFTNVDMPVQR, AAHLVPQGLLSESSR and VSTESLEPGSTEPAPQTDVAIGKPVR were identified.

(B) NanoLC-MS assays for specific peptide fragments of MSMEG_6160 protein. MSMEG_6160-specific peptide fragments with the amino acid sequences AAHALTESVAALR, AGAVKWLRLVLGALTK and AGYLAEDR were identified.

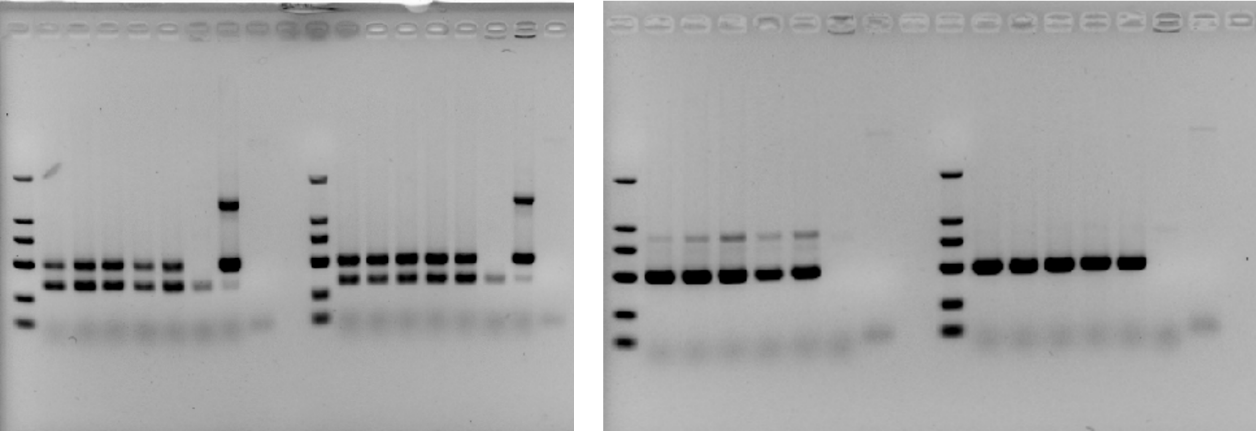

**Figure S7,** **related to Figure 5. Phage DNA circularization is disrupted by the Type III Druantia system.**

Uncropped gel Images, related to Figure 5B.

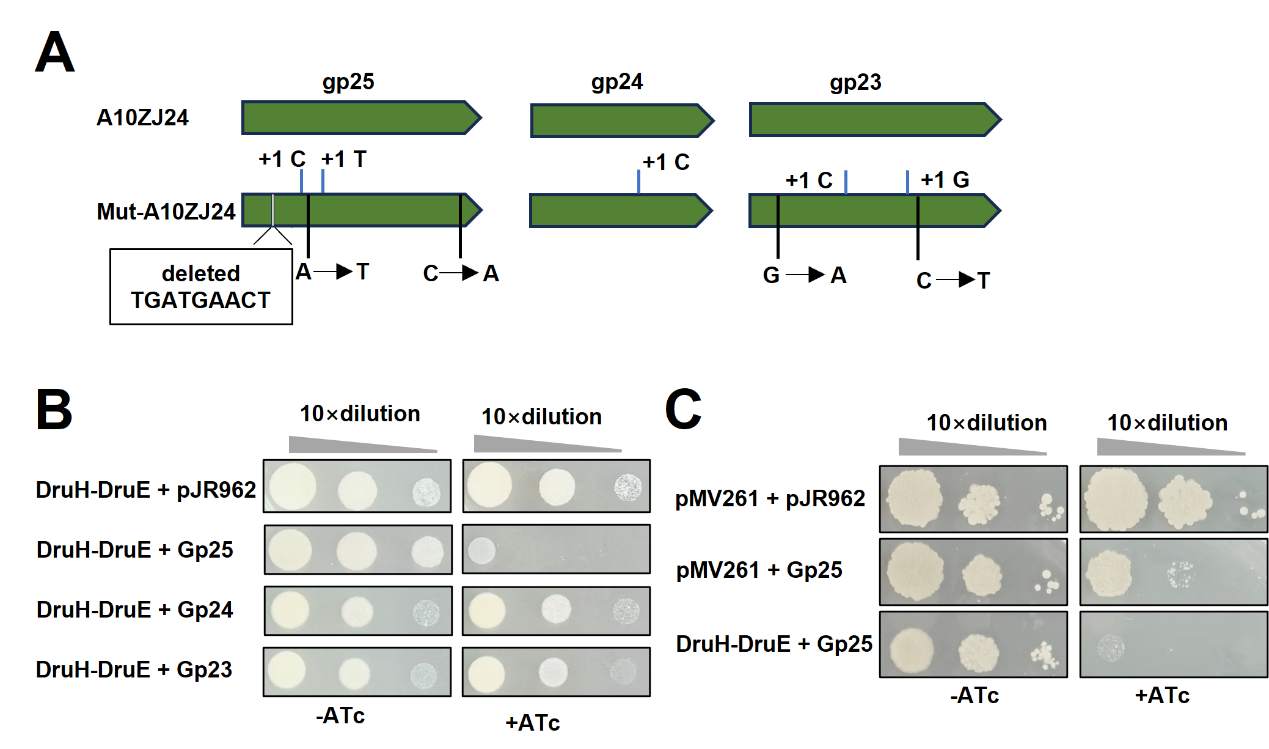

**Figure S8,** **related to Figure 6. Analysis of the mutated genes that confer phage to escape the type III Druantia defense system**

(A) Location of mutations found in phage that escape from the type III Druantia defense.

(B-C) Serial dilutions of cells were incubated in conditions with (+) or without (-) anhydrous tetracycline, bacterial viability was measured for strains in which the type III Druantia system was co-expressed with phage genes (*gp*23, *gp*24, *gp* 25).

pMV261+ pJR962 represents the wild-type strain. DruH-DruE + Gp25 represents the DruH-DruE strain with recombinant vector pJR962-*gp25*, in which Gp25 expression is induced by ATc. pMV261 + Gp25 represents the ATc-induced strain for Gp25 expression strain with an empty pMV261 vector. pMV261 + Gp25 represents the ATc-induced strain for gp25 expression strain with an empty pMV261 vector. DruH-DruE + Gp23 or 24 represents the DruH-DruE strain with recombinant vector pJR962-*gp23* or pJR962-*gp24*, in which Gp23 or Gp23 expression is induced by ATc.

**Supplementary Table 1. PCR primers used in this study.**

| Primers name | Sequence 5'-3' | Usage |
| --- | --- | --- |
| Ms1253-F | CGCGGTACCAGATCTTTAAATGTCTGCCCATTGCATTAGCCGTTG | Cloned to pMV261 |
| Ms1253-R | TCCAGCCCCCGATCCGACACTCAATACTGGTTGTCATTCCGCTTC | Cloned to pMV261 |
| Ms1253pro-F | CGCGGTACCAGATCTTTAAATGTCTGCCCATTGCATTAGCCGTTG | Cloned to pMV261 |
| Ms1253pro-R | GCATCAATACTGGTTGTCATTCCTTTTCCGTCCTCGACCCTTCGT | Cloned to pMV261 |
| Ms1254-F | GGGTCGAGGACGGAAAAGGAATGACAACCAGTATTGATGCAGTTG | Cloned to pMV261 |
| Ms1254-R | TTAACTACGTCGACATCGAT TCATGGAACAGCTTGCTGCTCGACC | Cloned to pMV261 |
| Ms1253-54-F | CGCGGTACCAGATCTTTAAA TGTCTGCCCATTGCATTAGCCGTTGT | Cloned to pMV261 |
| Ms1253-54-R | TTAACTACGTCGACATCGAT TCATGGAACAGCTTGCTGCTCGACC | Cloned to pMV261 |
| 1253-54(54K119A)-S1/F: | CGCGGTACCAGATCTTTAAATGTCTGCCCATTGCATTAGCCGTTG | Cloned to pMV261 |
| 1253-54(54K119A)-S1/R: | GCCTGACCCAGTTCCGGTGGCGACCA | Cloned to pMV261 |
| 1253-54(54K119A)-S2/F: | CCACCGGAACTGGGTCAGGCGCAACCGAATCGTTCCTCCTACCGA | Cloned to pMV261 |
| 1253-54(54K119A)-S2/R: | CCGATGTTGCAATGCACTGGATGGGACGGCCAGGAGCCACTCGGT | Cloned to pMV261 |
| 1253-54(54K119A)-S3/F: | CCAGTGCATTGCAACATCGGCCACT | Cloned to pMV261 |
| 1253-54(54K119A)-S3/R: | TTAACTACGTCGACATCGATTCATGGAACAGCTTGCTGCTCGACC | Cloned to pMV261 |
| 1253-54(54D255A)-S1/F | CGCGGTACCAGATCTTTAAATGTCTGCCCATTGCATTAGCCGTTG | Cloned to pMV261 |
| 1253-54(54D255A)-S1/R | GACGACGATGAATCGCCACTTGGAT | Cloned to pMV261 |
| 1253-54(54D255A)-S2/F | AGTGGCGATTCATCGTCGTCGCCGAAGCCCACGTCTACGACGGGA | Cloned to pMV261 |
| 1253-54(54D255A)-S2/R | CCACCATGGCTGCAAGTCCGTCTGCGGCTTCGGGTGCGCTGCCGA | Cloned to pMV261 |
| 1253-54(54D255A)-S3/F | CGGACTTGCAGCCATGGTGGAGCTC | Cloned to pMV261 |
| 1253-54(54D255A)-S3/R | TTAACTACGTCGACATCGATTCATGGAACAGCTTGCTGCTCGACC | Cloned to pMV261 |
| 1253-54(54A292D)-S1/F | CGCGGTACCAGATCTTTAAATGTCTGCCCATTGCATTAGCCGTTG | Cloned to pMV261 |
| 1253-54(54A292D)-S1/R | CGATGTTGCAATGCACTGGATGGGA | Cloned to pMV261 |
| 1253-54(54A292D)-S2/F | TCCAGTGCATTGCAACATCGGACACTGTCGGAAGTGATTCGGATC | Cloned to pMV261 |
| 1253-54(54A292D)-S2/R | GCCCCCTCGACGACAGGCAGGTGAATGCACCCTCGGTTGCCCTGA | Cloned to pMV261 |
| 1253-54(54A292D)-S3/F | CTGCCTGTCGTCGAGGGGGCCACAT | Cloned to pMV261 |
| 1253-54(54A292D)-S3/R | TTAACTACGTCGACATCGATTCATGGAACAGCTTGCTGCTCGACC | Cloned to pMV261 |
| 1253-54(54A1016D)-S1/F | CGCGGTACCAGATCTTTAAATGTCTGCCCATTGCATTAGCCGTTG | Cloned to pMV261 |
| 1253-54(54A1016D)-S1/R | ACGACCGGCACGCTGAAGGTAGTTG | Cloned to pMV261 |
| 1253-54(54A1016D)-S2/F | ACCTTCAGCGTGCCGGTCGTGATGGGCGCCGGTCAGGAGCCGCCG | Cloned to pMV261 |
| 1253-54(54A1016D)-S2/R | TCTGCCCAACCTTTTGATATCGCGCAGCGAGACCGTGTTTACCCT | Cloned to pMV261 |
| 1253-54(54A1016D)-S3/F | ATATCAAAAGGTTGGGCAGACACTG | Cloned to pMV261 |
| 1253-54(54A1016D)-S3/R | TTAACTACGTCGACATCGATTCATGGAACAGCTTGCTGCTCGACC | Cloned to pMV261 |
| 1253-54(54C1516A)-S1/F | CGCGGTACCAGATCTTTAAATGTCTGCCCATTGCATTAGCCGTTG | Cloned to pMV261 |
| 1253-54(54C1516A)-S1/R | CTCACTCACCCGTTTGAGAGCGCAT | Cloned to pMV261 |
| 1253-54(54C1516A)-S2/F | CTCTCAAACGGGTGAGTGAGGCTGAGTGCGGGCCGGAAACGTCGT | Cloned to pMV261 |
| 1253-54(54C1516A)-S2/R | TTAACTACGTCGACATCGATTCATGGAACAGCTTGCTGCTCGACC | Cloned to pMV261 |
| (53R536A)-S1/F | CGCGGTACCAGATCTTTAAATGTCTGCCCATTGCATTAGCCGTTG | Cloned to pMV261 |
| (53R536A)-S1/R | caaattggagtaaaggtcgccgCaagcccctcgactacagtgaat | Cloned to pMV261 |
| (53R536A)-S2/F | gcgacctttactccaatttggcgacgt | Cloned to pMV261 |
| (53R536A)-S2/R | TTAACTACGTCGACATCGATTCATGGAACAGCTTGCTGCTCGACC | Cloned to pMV261 |
| (53D240A)-S1/R | ctccatcctgatgccaccgcgGcgagcagtGcaaacgaacgatcggccacgtcg | Cloned to pMV261 |
| (53D240A)-S2/F | gcggtggcatcaggatggagcgtcg | Cloned to pMV261 |
| (53E842A)-S1/R | gccctaacggtccagagccgtGcgaaatcaatcacctcgacaggtt | Cloned to pMV261 |
| (53E842A)-S2/F | cggctctggaccgttagggcgctgct | Cloned to pMV261 |
| (53R864/865A)-S1/R | tagacctcagtatcgatagccGctGcgactGcccgaatgcgctgtccaagact | Cloned to pMV261 |
| (53R864/865A)-S2/F | gctatcgatactgaggtctacgcaaatcc | Cloned to pMV261 |
| (53E1096/1099A)-S1/R | gtgaccaactgaggaacaaccGcagccaacGccgcccaggctcgtttctctcga | Cloned to pMV261 |
| (53E1096/1099A)-S2/F | gttgttcctcagttggtcaccatcgac | Cloned to pMV261 |
| (53D1108A)-S1/R | atcagctcggcgataatcaagGcgatggtgaccaactgaggaacaa | Cloned to pMV261 |
| (53D1108A)-S2/F | ttgattatcgccgagctgatcatcgg | Cloned to pMV261 |
| (53E998A)-S1/R | accgcgaggatgcgggacgacGcggaaagcaagccttgaggcacaa | Cloned to pMV261 |
| (53E998A)-S2/F | tcgtcccgcatcctcgcggtgat | Cloned to pMV261 |
| (53 R410A)-S1/R | ATCTGGAACGATCCGCTGACGGCCAGCGCGTCGCG | Cloned to pMV261 |
| (53 R410A)-S2/F | gtcagcggatcgttccagatcatcg | Cloned to pMV261 |
| (53 E651A) -S1/R | ACACCCACGTCGAGGACAAGTGCTCCAGGCCGCTC | Cloned to pMV261 |
| (53 E651A) -S2A/F | cttgtcctcgacgtgggtgtggtag | Cloned to pMV261 |
| (53 T675A) -S1/R | CGCCCGCTCTTCGGCGAAAGGGCCTGAATCGACCG | Cloned to pMV261 |
| (53 T675A) -S2/F | ctttcgccgaagagcgggcgtatcg | Cloned to pMV261 |
| (53 T696A) -S1/R | ACGTGCGGATACTCACCCAAAGCGTCTGCAATCTC | Cloned to pMV261 |
| (53 T696A) -S2/F | ttgggtgagtatccgcacgtcagcc | Cloned to pMV261 |
| (54 E543A) -S1/R | GTCACCGTCGTCGACGGCGGCGGCT | Cloned to pMV261 |
| (54 E543A) -S2/F | ccgccgtcgacgacggtgac gCagtctctggcgat | Cloned to pMV261 |
| (54 R581A) -S1/R | GAGTTTGCGCACTGGGCGGAGCTCG | Cloned to pMV261 |
| (54 R581A) -S2/F | tccgcccagtgcgcaaactc GCgcagcgtggcgag | Cloned to pMV261 |
| (54 R823A) -S1/R | GCCCGGCAGCCATGAGAGGACCTTT | Cloned to pMV261 |
| (54 R823A) -S2/F | tcctctcatggctgccgggc GCtggaacgaatcgc | Cloned to pMV261 |
| (54 V857A) -S1/R | AAGGAACTCCCATACGCCCTTCAAG | Cloned to pMV261 |
| (54 V857A) -S2/F | agggcgtatgggagttcctt gCgcgcagagaaact | Cloned to pMV261 |
| (54 R859A) -S1/R | GCGCACAAGGAACTCCCATACGCCC | Cloned to pMV261 |
| (54 R859A) -S2/F | tatgggagttccttgtgcgc GCagaaactccggtg | Cloned to pMV261 |
| (54 V1615A) -S1/R | TCCGTTGGCGGCCTCGAATCCGAGT | Cloned to pMV261 |
| (54 V1615A) -S2/F | gattcgaggccgccaacgga gCtcctatccctatt | Cloned to pMV261 |
| Lysogen-F | CGTGATAGTCCAGACCGGCA | PCR |
| Lysogen-R | TTCGTGTCCCCCTGGACCCA | PCR |
| bacterial-F | ATGGCAAAGAAAGTGACCGT | PCR |
| bacterial-R | CTAAGTTGCCGCGTGGAATG | PCR |
| Phage-F | ATGGAAGACGACGGGACCAA | PCR |
| Phage-R | CTACGGCTCCACGGGTGGGA | PCR |
| Circular-F | CAGAACGGGCTGCCGGTAGT | PCR |
| Circular-R | GCGATCCGCTGTTGAGTTGT | PCR |
| pTRG-Ms1253-F | GCGGCCGC GTGTCGTTATGTCCCGGTTTACCGC | Cloned to pTRG |
| pTRG-Ms1253-R | TCTAGA TCAATACTGGTTGTCATTCCGCTTC | Cloned to pTRG |
| (53D410A)-S1/R | ATCTGGAACGATCCGCTGACGGCCAGCGCGTCGCG | Cloned to pTRG |
| (53D410A)-S2/F | gtcagcggatcgttccagatcatcg | Cloned to pTRG |
| (53 E651A) -S1/R | ACACCCACGTCGAGGACAAGTGCTCCAGGCCGCTC | Cloned to pTRG |
| (53 E651A) -S2A/F | cttgtcctcgacgtgggtgtggtag | Cloned to pTRG |
| pBT-Ms1254-F | GGCCTGAAGAGACGTTTGGCATGACAACCAGTATTGATGCAGTTG | Cloned to pBT |
| pBT-Ms1254-R | ACGCACGCCCGAGATGATACTCATGGAACAGCTTGCTGCTCGACC | Cloned to pBT |
| (54 R859A) -S1/R | GCGCACAAGGAACTCCCATACGCCC | Cloned to pBT |
| (54 R859A) -S2/F | tatgggagttccttgtgcgc GCagaaactccggtg | Cloned to pBT |
| (54 R823A) -S1/R | GCCCGGCAGCCATGAGAGGACCTTT | Cloned to pBT |
| (54 R823A) -S2/F | tcctctcatggctgccgggc GCtggaacgaatcgc | Cloned to pBT |
| COS-F | CTCTGGTATCGCGGATAGGG | qRT-PCR |
| COS-R | GGGTATCGCTGCAGGTAGG | qRT-PCR |
| 16SrRNART-F | GATACGGGCAGACTAGAGTA | qRT-PCR |
| 16SrRNART-R | GGGTATCTAATCCTGTTCGC | qRT-PCR |
| Dut-Ms1253-F: | ACCATCATCACCACAGCCAGGTGTCGTTATGTCCCGGTTTACCGC | Cloned to pRSF-Dut |
| Dut-Ms1253-R: | CTTAAGCATTATGCGGCCGCTCAATACTGGTTGTCATTCCGCTTC | Cloned to pRSF-Dut |
| Gp48-F: | TCAGTGATAGAGAAGGCGGTATGAGTACCAGCACCATCACGACGC | Cloned to pLJR-962 |
| Gp48-R： | GCCTTCTGCTTAGCTAATCATTAGAGGAGTCCGATCTGACCCATG | Cloned to pLJR-962 |
| Gp49-F: | TCAGTGATAGAGAAGGCGGTGTGGACACCCTCGTACAGCTTGAGG | Cloned to pLJR-962 |
| Gp49-R： | GCCTTCTGCTTAGCTAATCATCATTCGAGCCCCCAGGGCCTCGTC | Cloned to pLJR-962 |
| Gp50-F: | TCAGTGATAGAGAAGGCGGTATGAAGGGCTTCCGGGAGGAGATCG | Cloned to pLJR-962 |
| Gp50-R： | GCCTTCTGCTTAGCTAATCATTACCTCCTATTGAATTGCAACGCC | Cloned to pLJR-962 |
| Mra_3684-F: | CGCGGTACCAGATCTTTAAAATACATCCGCGGAACCGTCTTCGGG | Cloned to pMV261 |
| Mra_3684-R: | TTAACTACGTCGACATCGATTCACGGTGATTCCTCACTTAACTCG | Cloned to pMV261 |
| Ms6160-F: | CGCGGTACCAGATCTTTAAAAGCCGAAGCCCTTCTCCGCGTTGAA | Cloned to pMV261 |
| Ms6160-R: | TTAACTACGTCGACATCGATCTACTTCGTCAGCGCGCCGAGCACC | Cloned to pMV261 |
| 53-Mra_3684-F1: | CGCGGTACCAGATCTTTAAATGTCTGCCCATTGCATTAGCCGTTG | Cloned to pMV261 |
| 53-Mra_3684-F1: | TGGCTGCCGAAACTCGCCATTCAATACTGGTTGTCATTCCGCTTC | Cloned to pMV261 |
| 53-Mra_3684-F2: | GGAATGACAACCAGTATTGAATGGCGAGTTTCGGCAGCCACCTGC | Cloned to pMV261 |
| 53-Mra_3684-R2: | TTAACTACGTCGACATCGATTCACGGTGATTCCTCACTTAACTCG | Cloned to pMV261 |
| 53-Ms6160-F1: | CGCGGTACCAGATCTTTAAATGTCTGCCCATTGCATTAGCCGTTG | Cloned to pMV261 |
| 53- Ms6160-F1: | TCCAGCCCCCGATCCGACACTCAATACTGGTTGTCATTCCGCTTC | Cloned to pMV261 |
| 53- Ms6160-F2: | GGAATGACAACCAGTATTGAGTGTCGGATCGGGGGCTGGAATTCG | Cloned to pMV261 |
| 53- Ms6160-R2: | TTAACTACGTCGACATCGATCTACTTCGTCAGCGCGCCGAGCACC | Cloned to pMV261 |

**Supplementary Table 2. Strains, phages and plasmids used in this study.**

| **Plasmid, phage or Strain** | **Relevant genotype or feature** | | **Source or reference** |
| --- | --- | --- | --- |
| Plasmid |  | |  |
| pRSF-Dut | *Kan^r^*, T7 lac promoter, N-terminal His_6_ | | Novagen |
| pRSF -*MSMEG_5860* | *MSMEG_5860* inserted in *BamH* I*-Hind* Ⅲ of pRSF-Dut | | This study |
| pJR962 | including Sth1 sgRNA scaffold and dCas9, *Hyg^r^* instead of  *Kan^r^*) This study | | Previous study ^[S1]^ |
| pJR962-*A10ZJ24gp48* | *A10ZJ24gp48* inserted in *Cla* I*-Not* Ⅰof pLJR962 | | This study |
| pJR962- *A10ZJ24gp49* | *A10ZJ24gp49* inserted in *Cla* I*-Not* Ⅰ of pLJR962 | | This study |
| pJR962- *A10ZJ24gp50* | *A10ZJ24gp50* inserted in *Cla* I*-Not* Ⅰ of pLJR962 | | This study |
| pJR962-*Mut-gp48* | *Mut-gp48* inserted in *Cla* I*-Not* Ⅰ of pLJR962 | | This study |
| pJR962-*MSMEG_1253* | *MSMEG_1253* inserted in *Cla* I*-Not* Ⅰ of pLJR962 | | This study |
| pMV261 | *Kan^r^*, pAL5000 replicon | | Previous study^[S1]^ |
| pMV261-*MSMEG_1253*-1254 | *MSMEG_1253*-1254 inserted in *Xba* I-*Hind* Ⅲ sites of pMV261 | | This study |
| pMV261-*MSMEG_1253* | *MSMEG_1253* inserted in *Xba* I-*Hind* Ⅲ sites of pMV261 | | This study |
| pMV261-*MSMEG_1254* | *MSMEG_1254* inserted in *Xba* I-*Hind* Ⅲ sites of pMV261 | | This study |
| pMV261-*MSMEG_6160* | *MSMEG_6160* inserted *in Xba* I-*Hind* Ⅲ *of* pMV261 | | This study |
| pMV261-*Mra_3684* | *Mra_3684 inserted in Xba* I-*Hind* Ⅲ *of* pMV261 | | This study |
| pMV261-*DruH/MSMEG_6160* | *MSMEG_1253/MSMEG_6160 inserted in Xba* I-*Hind* Ⅲ *of* pMV261 | | This study |
| pMV261-*DruH/Mra_3684* | *MSMEG_1253/Mra_3684 inserted in Xba* I-*Hind* Ⅲ *of* pMV261 | | This study |
| pBT | Bacterial 2-hybrid assay bait vector | | Stratagene |
| pTRG | Bacterial 2-hybrid assay bait vector | | Stratagene |
| pBT-*MSMEG_1254* | *MSMEG_1254*in *Not* I-*Xba* Ⅰ sites of pBT | | This study |
| pTRG-*MSMEG_1253* | *MSMEG_1253* in *Not* I-*Xba* Ⅰ sites of pTRG | | This study |
| Strain |  | |  |
| *E. coli* BL21 (DE3) |  | | TaKaRa |
| *M. smegmatis* MC^2^ 155 |  | | ATCC |
| *M. tuberculosis* H37Ra |  | | ATCC |
| WT*/*pMV261 | | *M. smegmatis* MC^2^ 155 with pMV261 | This study |
| DruH-DruE | *M. smegmatis* MC^2^ 155 with pMV261-*MSMEG_1253-1254* | | This study |
| DruE | *M. smegmatis* MC^2^ 155 with pMV261-*MSMEG_1254* | | This study |
| DruH | *M. smegmatis* MC^2^ 155 with pMV261-*MSMEG_1253* | | This study |
| DruH^E1096/1099A^-DruE | *M. smegmatis* MC^2^ 155 with pMV261-*MSMEG_1253(*E1096/1099A*)-1254* | | This study |
| DruH^E842A^-DruE | *M. smegmatis* MC^2^ 155 with pMV261-  *MSMEG_1253*(E842A)*-1254* | | This study |
| DruH^R536A^-DruE | *M. smegmatis* MC^2^ 155 with pMV261-  *MSMEG_1253*(R536A)*-1254* | |  |
| DruH^D240A^-DruE | *M. smegmatis* MC^2^ 155 with pMV261-  *MSMEG_1253*(D240A)*-1254* | |  |
| **Continued** | |  |  |
| **Plasmid, phage or Strain** | | **Relevant genotype or feature** | **Source or reference** |
| DruH^R864/865A^-DruE | *M. smegmatis* MC^2^ 155 with pMV261-*MSMEG_1253*(R864/865A)*-1254* | | This study |
| DruH^D1108A^-DruE | *M. smegmatis* MC^2^ 155 with pMV261-*MSMEG_1253*(D1108A)*-1254* | | This study |
| DruH^E998A^-DruE | *M. smegmatis* MC^2^ 155 with pMV261-  *MSMEG_1253*(E998A)*-1254* | | This study |
| DruH-DruE^K119A^ | *M. smegmatis* MC^2^ 155 with pMV261-  *MSMEG_1253-1254*(K119A) | | This study |
| DruH-DruE^D225A^ | *M. smegmatis* MC^2^ 155 with pMV261-  *MSMEG_1253-1254*(D225A) | | This study |
| DruH-DruE^A292D^ | *M. smegmatis* MC^2^ 155 with pMV261-  *MSMEG_1253-1254*(A292D) | | This study |
| DruH-DruE^A1016D^ | *M. smegmatis* MC^2^ 155 with pMV261-  *MSMEG_1253-1254*(A1016D) | | This study |
| DruH-DruE^C1516A^ | *M. smegmatis* MC^2^ 155 with pMV261-  *MSMEG_1253-1254*(C1516A) | | This study |
| DruH-DruE^P800A^ | *M. smegmatis* MC^2^ 155 with pMV261-  *MSMEG_1253-1254*(P800A) | | This study |
| DruH^R410A^-DruE | *M. smegmatis* MC^2^ 155 with pMV261-  *MSMEG_1253* (R410A) *-1254* | | This study |
| DruH^E651A^-DruE | *M. smegmatis* MC^2^ 155 with pMV261-  *MSMEG_1253* (E651A) *-1254* | | This study |
| DruH^T675A^-DruE | *M. smegmatis* MC^2^ 155 with pMV261-  *MSMEG_1253* (T675A) *-1254* | | This study |
| DruH^T696A^-DruE | *M. smegmatis* MC^2^ 155 with pMV261-  *MSMEG_1253* (T696A) *-1254* | | This study |
| DruH-DruE^E543A^ | *M. smegmatis* MC^2^ 155 with pMV261-  *MSMEG_1253-1254*(E543A) | | This study |
| DruH-DruE^R581A^ | *M. smegmatis* MC^2^ 155 with pMV261-  *MSMEG_1253-1254*(R581A) | | This study |
| DruH-DruE^R823A^ | *M. smegmatis* MC^2^ 155 with pMV261-  *MSMEG_1253-1254*(R823A) | | This study |
| DruH-DruE^V857A^ | *M. smegmatis* MC^2^ 155 with pMV261-  *MSMEG_1253-1254*(V857A) | | This study |
| DruH-DruE^R859A^ | *M. smegmatis* MC^2^ 155 with pMV261-  *MSMEG_1253-1254*(R859A) | | This study |
| DruH-DruE V1615A | *M. smegmatis* MC^2^ 155 with pMV261-  *MSMEG_1253-1254*(V1615A) | | This study |
| pMV261 + Gp25 | *M. smegmatis* MC^2^ 155 with pMV261 and  pJR962-*A10ZJ24gp25* | | This study |
| ***Continued*** | |  |  |
| **Plasmid, phage or Strain** | | **Relevant genotype or feature** | **Source or reference** |
| DruH-DruE + Gp25 | | *M. smegmatis* MC^2^ 155 with pMV261-  *MSMEG_1253-1254* and pJR962-*A10ZJ24gp25* | This study |
| DruH-DruE^EA292D^ + Gp25 | | *M. smegmatis* MC^2^ 155 with pMV261-  *MSMEG_1253-1254* (A292D) and pJR962-*A10ZJ24gp25* | This study |
| DruH-DruE^A1016D^ + Gp25 | | *M. smegmatis* MC^2^ 155 with pMV261-  *MSMEG_1253-1254* (A1016D) and pJR962-A10ZJ24*gp25* | This study |
| pMV261 + Mut-Gp25 | | *M. smegmatis* MC^2^ 155 with pMV261 and pJR962-Mut-*gp25* | This study |
| DruH-DruE + Mut-Gp25 | | *M. smegmatis* MC^2^ 155 with pMV261-  *MSMEG_1253-1254* and pJR962-Mut-*gp25* | This study |
| Mra/pMV261 | | *M. tuberculosis* H37Ra with pMV261 | This study |
| (Mra_3684) | | *M. tuberculosis* H37Rv with pMV261- *Mra_3684* | This study |
| (DruH-Mra_3684) | | *M. tuberculosis* H37Rv with pMV261-DruH/*Mra_3684* | This study |
| (DruH-DruE) | | *M. tuberculosis* H37Rv with pMV261-*MSMEG_1253-1254* | This study |
| (DruH-DruE^K119A^) | | *M. tuberculosis* H37Rv with pMV261- *MSMEG_1253-1254* (K119A) | This study |
| (DruH-DruE^D225A^) | | *M. tuberculosis* H37Rv with pMV261- *MSMEG_1253-1254* (D225A) | This study |
| (DruH-DruE^A292D^) | | *M. tuberculosis* H37Ra with pMV261- *MSMEG_1253-1254* (A292D) | This study |
| (DruH-DruE^A1016D^) | | *M. tuberculosis* H37Ra with pMV261- *MSMEG_1253-1254* (A1016D) | This study |
| DruH/DruE | | E. coli reporter strain with pBT-*MSMEG_1254* and pTRG-*MSMEG_1253* | This study |
| DruH/DruE^R823A^ | | E. coli reporter strain with pBT-*MSMEG_1254*(R823A) and pTRG-*MSMEG_1253* | This study |
| DruH/DruE^R859A^ | | E. coli reporter strain with pBT-*MSMEG_1254*(R859A) and pTRG-*MSMEG_1253* | This study |
| DruH^R410A^/DruE^R823A^ | | E. coli reporter strain with pBT-*MSMEG_1254*(R823A) and pTRG-*MSMEG_1253*(R410A) | This study |
| DruH^E651A^/DruE^R823A^ | | E. coli reporter strain with pBT-*MSMEG_1254*(R823A) and pTRG-*MSMEG_1253*(E651A) | This study |
| A1ZJ29 | |  | This study |
| F1GX13 | |  | This study |
| A4ZJ24 | |  | This study |
| A22GX2 | |  | This study |
| A9GX2 | |  | This study |
| A1GX133 | |  | This study |
| Plasmid, phage or Phage | | Relevant genotype or feature | Source or reference |
| A1GX133 | |  | This study |
| A1GX135 | |  | This study |
| A1GX136 | |  | This study |
| A1GX130 | |  | This study |
| A9GX19 | |  | This study |
| A2GX14 | |  | This study |
| A3HA12 | |  | This study |
| A3JX14 | |  | This study |
| K5GX11 | |  | This study |
| K5HA9 | |  | This study |
| K5HA7 | |  | This study |
| K5GX12 | |  | This study |
| K5GX5 | |  | This study |
| K4JX5 | |  | This study |
| K4GX10 | |  | This study |
| K4SN7 | |  | This study |
| K4HE12 | |  | This study |
| K4JX17 | |  | This study |
| L2JS15 | |  | This study |
| L2GX4 | |  | This study |
| L2GX23 | |  | This study |
| L2HB19 | |  | This study |
| L2JX21 | |  | This study |
| A1SD115 | |  | This study |

**References**

Li, X. et al. Mycobacterial phage TM4 requires a eukaryotic-like Ser/Thr protein kinase to silence and escape anti-phage immunity. *Cell Host Microbe* **31**, 1469-1480
